# Retinal network dysfunction precedes structural degeneration in severe *GUCA1A* cone-rod dystrophy

**DOI:** 10.64898/2026.06.23.734025

**Authors:** Anna Avesani, Giuditta Dal Cortivo, Sabrina Asteriti, Giorgia Targa, Laura Veschetti, Valerio Marino, Amedeo Biasi, Carmen Longo, Barbara Cisterna, Karolina Saran, Giovanni Malerba, Andrzej T. Foik, Marco Cambiaghi, Lorenzo Cangiano, Daniele Dell’Orco

## Abstract

Autosomal dominant cone-rod dystrophy caused by *GUCA1A* mutations is generally viewed as a disorder of phototransduction, yet the mechanisms linking photoreceptor dysfunction to progressive vision loss remain unclear. Here, using a knock-in mouse carrying the severe GCAP1 p.(E111V) variant, we show that retinal network dysfunction precedes structural degeneration. Mutant mice exhibited delayed rod photoresponses, increased light sensitivity, selective visuospatial deficits, and progressive impairment of visually evoked responses in the superior colliculus and visual cortex, demonstrating propagation of functional deficits beyond photoreceptors. Transcriptomic and ultrastructural analyses revealed early synaptic, mitochondrial and inflammatory alterations despite largely preserved retinal architecture. Acute ex vivo delivery of recombinant wild-type GCAP1 partially restored mutant rod photoresponse kinetics, indicating that these early functional deficits remain biochemically modifiable. These findings redefine severe *GUCA1A*-associated disease as a progressive disorder of retinal network function, identifying an early therapeutic window before structural degeneration.

**One-Sentence Summary:** Visual function breaks down long before photoreceptors are lost, opening an early window for intervention.

## Introduction

Cone-rod dystrophies (CORDs) are inherited retinal dystrophies characterized by primary cone dysfunction followed by progressive visual loss and secondary rod degeneration, affecting approximately 1 in 40,000 individuals worldwide (*1*). Although the genetic architecture of CORDs spans dozens of loci (*2, 3*), the mechanisms by which an initially photoreceptor-confined molecular defect propagates through retinal circuits to impair visual function remain poorly understood. This question is clinically relevant because CORDs cause early-onset photophobia, color vision defects, visual acuity loss and progressive central scotoma, and no disease-modifying therapy is currently approved. Defining the earliest events linking photoreceptor dysfunction to vision loss is therefore essential for identifying effective therapeutic strategies in inherited retinal degeneration.

Guanylate cyclase-activating protein 1 (GCAP1), encoded by the *GUCA1A* gene, is a neuronal Ca²⁺ sensor that regulates retinal guanylate cyclase 1 (GC1) during response recovery and light adaptation in both rods and cones (*4*). Light-induced reductions in intracellular Ca²⁺ trigger the transition of GCAP1 from its inhibitory Ca²⁺-bound state to an activating Mg²⁺-bound form, thereby stimulating GC1 to restore cGMP levels and the circulating current (*5–7*). Consistent with its higher expression in cones than rods (*8, 9*), *GUCA1A* mutations predominantly affect cone-mediated vision. To date, more than 20 missense variants have been associated with autosomal dominant cone dystrophy (COD), cone-rod dystrophy (CORD) and, less frequently, retinitis pigmentosa (RP) (*10–12*), including several that have been extensively characterized biochemically and clinically (*13–18*). Despite their clinical heterogeneity, all well-characterized disease-causing variants share a common pathogenic mechanism: impaired Ca²⁺-dependent regulation of GC1 by mutant GCAP1, resulting in constitutive cyclase activation and toxic accumulation of cGMP and Ca²⁺ in photoreceptors (*19*). Whether disruption of this photoreceptor-specific signaling pathway remains confined to photoreceptors or extends beyond them to impair retinal network function is unknown.

Addressing this question requires animal models that faithfully reproduce the molecular defect under physiological conditions. Three mouse models have previously been developed to investigate *GUCA1A*-associated adCORD. Olshevskaya et al. generated transgenic mice overexpressing bovine Y99C-GCAP1, which developed elevated intracellular Ca²⁺ and progressive photoreceptor degeneration in a transgene dose-dependent manner, resembling retinitis pigmentosa rather than CORD (*20*). Buch et al. introduced the E155G substitution into the endogenous *GUCA1A* locus, recapitulating late-onset cone-predominant degeneration and cGMP accumulation (*21*). Jiang et al. generated transgenic mice expressing L151F-GCAP1, in which meaningful structural and functional deterioration became evident only after approximately nine months of age (*22*). Although these models have provided important mechanistic insights, none combines allelic fidelity at the endogenous locus, physiological expression under native regulatory control, and a mutation associated with the most severe biochemical and clinical phenotype. These limitations have hindered investigation of the earliest functional consequences of severe *GUCA1A* mutations and motivated the development of a physiologically faithful knock-in model.

Among all disease-causing *GUCA1A* variants, we previously identified the p.(E111V) substitution in GCAP1 as the mutation associated with the most severe molecular phenotype, accompanied by an unusually severe clinical presentation (*16*). E111V disrupts Ca²⁺ coordination at the high-affinity EF-hand 3, reducing the apparent Ca²⁺ affinity of GCAP1 by approximately 80-fold and causing constitutive activation of GC1 well beyond the physiological Ca²⁺ range. Clinically, affected individuals present with early-onset nystagmus, photophobia and progressive central outer retinal atrophy, with virtually absent scotopic electroretinogram responses by the third decade of life. The molecular consequences of this profound disruption of Ca²⁺ and cGMP homeostasis have been investigated extensively in vitro, providing mechanistic insight (*23*) and establishing proof-of-concept for biochemical intervention using recombinant wild-type GCAP1 (*24, 25*). These observations identify E111V as an ideal variant with which to investigate how severe disruption of phototransduction impacts retinal function beyond the photoreceptor.

To investigate how severe dysregulation of GCAP1 signaling affects retinal function in vivo, we generated a CRISPR/Cas9 knock-in mouse carrying the E111V mutation at the endogenous *Guca1a* locus. Using complementary structural, functional, behavioral and transcriptomic approaches, we investigated how severe disruption of phototransduction influences retinal and visual pathway function. Our findings reveal an unexpected stage of disease progression in which retinal and visual network dysfunction precedes overt structural degeneration and identify an early opportunity for biochemical intervention.

## RESULTS

### Retinal architecture remains largely preserved during the early stages of severe *GUCA1A* disease

Because our central hypothesis predicts that functional abnormalities emerge early during disease progression, we first asked whether the severe E111V mutation causes detectable structural alterations in the retina. To this end, we generated a CRISPR/Cas9 knock-in mouse carrying the endogenous *Guca1a* p.(E111V) mutation and assessed retinal morphology longitudinally at postnatal days P30, P120 and P270, corresponding to successive stages of disease progression in affected patients.

Retinal architecture remained remarkably preserved throughout the early stages of disease. Quantitative analysis of outer nuclear layer (ONL) thickness (**Fig. 1A, B**), normalized to WT at P30 (fold change), revealed no significant differences among genotypes at either P30 (WT: 1.00 ± 0.11; E111V⁺/⁻: 1.02 ± 0.03; E111V⁺/⁺: 0.93 ± 0.09) or P120 (E111V⁺/⁻: 1.00 ± 0.11; E111V⁺/⁺: 0.97 ± 0.07). A significant reduction in ONL thickness became apparent only at P270 in E111V⁺/⁺ mice (0.74 ± 0.12) relative to age-matched WT retinas (1.00 ± 0.04), whereas heterozygous animals remained indistinguishable from controls (1.07 ± 0.15). No significant differences were detected in inner nuclear layer (INL) thickness at any age (**Fig. 1B**). Because homozygous mice consistently exhibited the most pronounced phenotype, subsequent mechanistic analyses focused on comparisons between E111V⁺/⁺ and WT animals.

**Fig. 1.**
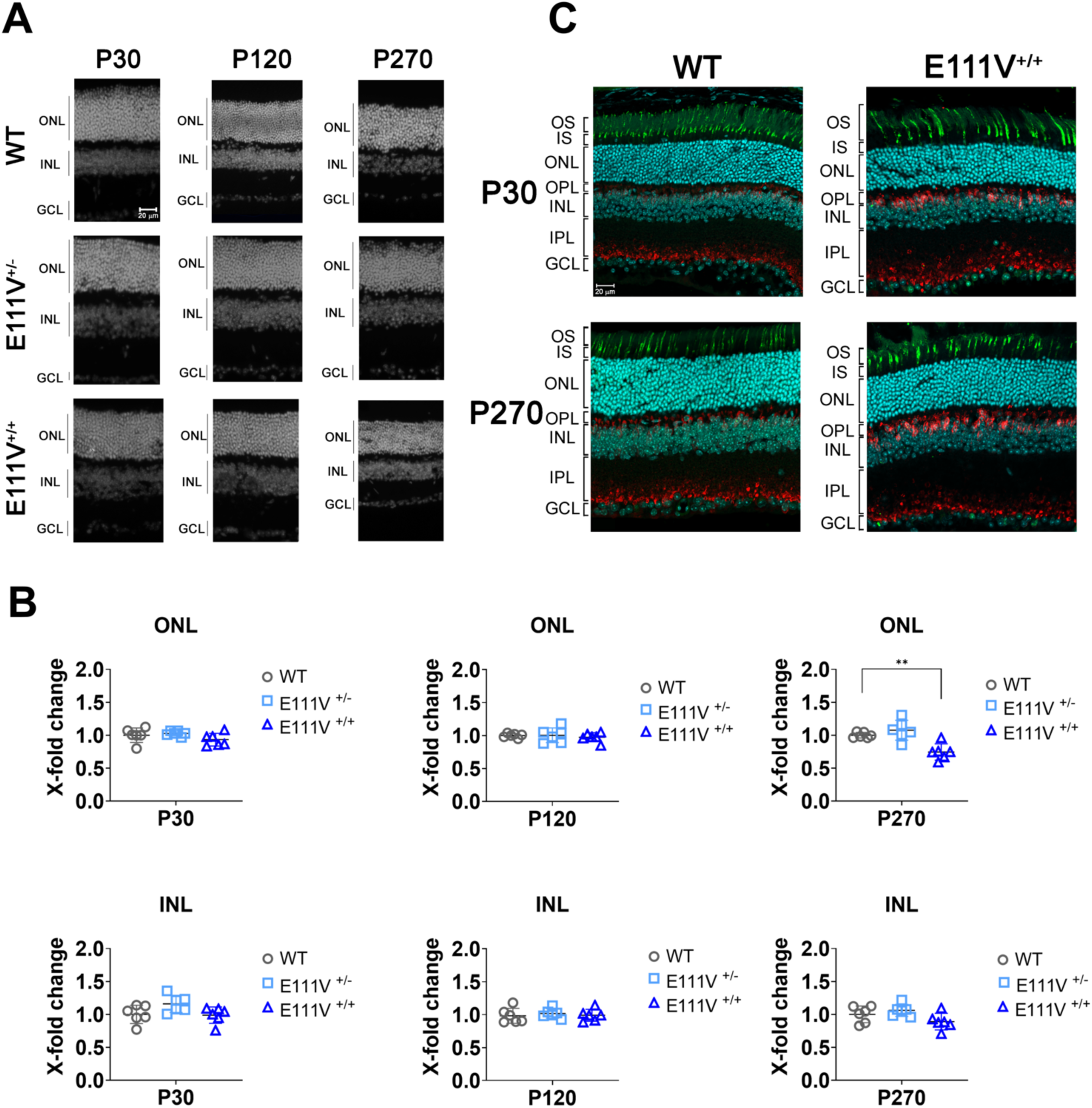
Characterization of retinal degeneration and ONL loss in E111V mice. (**A**) Representative DAPI immunofluorescence images from n = 6 mice per genotype at P30, P120 and P270. Images acquired with Olympus microscope (20× objective; scale bar: 20 µm). (**B**) ONL and INL thickness normalized to WT (X-fold change). Data analyzed by ordinary one-way ANOVA with multiple comparisons (GraphPad Prism). Significant differences: *p < 0.05, **p < 0.01, ***p < 0.001. (**C**) Immunofluorescence labeling of cone photoreceptors (PNA-488nm) and rod bipolar cells (PKCα, 647 nm) in retinal cryosections from WT and E111V^+/+^ mice at P30 and P270. Shown are z-stack projections of 5 optical sections (0.7 µm z-spacing). Images acquired on Fluoview 4000 confocal microscope (20× objective; scale bar, 20 µm).

Histological examination of hematoxylin and eosin-stained sections confirmed that retinal lamination remained largely preserved throughout the observation period (**Supplementary Fig. S1**). To determine whether subtle cellular alterations accompanied the limited ONL thinning, we next examined cone photoreceptors and rod bipolar cells by immunofluorescence (**Fig. 1C**). At P30, peanut agglutinin (PNA) labeling revealed no overt cone loss in E111V⁺/⁺ retinas, although staining intensity appeared modestly reduced relative to WT. Rod bipolar cells, identified by protein kinase Cα (PKCα), displayed preserved laminar organization across both the outer and inner plexiform layers, with no evidence of disrupted synaptic stratification. Longitudinal analysis revealed the expected age-dependent decline in PNA labeling in both genotypes, whereas PKCα immunoreactivity remained largely stable but showed increased labeling within the outer plexiform layer of E111V⁺/⁺ retinas at later ages, consistent with subtle genotype-dependent changes in bipolar cell organization. Notably, despite the severe retinal phenotype observed in patients carrying the E111V mutation, mutant mice did not exhibit substantial cone loss during the period examined.

Together, these observations indicate that severe E111V-GCAP1 expression produces only limited structural alterations during the early stages of disease progression, suggesting that overt retinal degeneration is unlikely to account for the early functional abnormalities investigated in the following sections.

### Rod photoresponses are altered before overt retinal degeneration in E111V mice

The limited structural alterations observed during early disease progression prompted us to determine whether retinal function was already compromised. We therefore examined rod photoreceptor responses by ex vivo electroretinography (ERG) using pharmacologically isolated scotopic a-waves from dark-adapted retinas (**Fig. 2**). Despite the largely preserved retinal architecture described above, E111V⁺/⁺ rods exhibited robust functional abnormalities that were already evident in young adult animals. Mutant rods were consistently more light sensitive than WT controls, as indicated by a significant reduction in the half-saturating flash strength (i50) both in young adults (P74; 19.2 ± 2.1 vs 26.2 ± 1.9 photons/µm²; P < 0.001; n = 10 E111V⁺/⁺, 10 WT) and in aged animals (P378; 24.6 ± 1.2 vs 34.9 ± 1.0 photons/µm²; P < 0.001; n = 27 E111V⁺/⁺, 25 WT) (**Fig. 2A,B**). Response kinetics (TTP@i50) were likewise significantly slowed in mutant rods at both ages, with prolonged time-to-peak values (young adults: 158 ± 9 vs 110 ± 3 ms; P < 0.001; aged animals: 145 ± 3 vs 101 ± 1 ms; P < 0.001). A significant reduction in saturating response amplitude was detected only in older mutant mice (P < 0.001), indicating relatively slow progression of photoreceptor dysfunction.

**Fig. 2.**
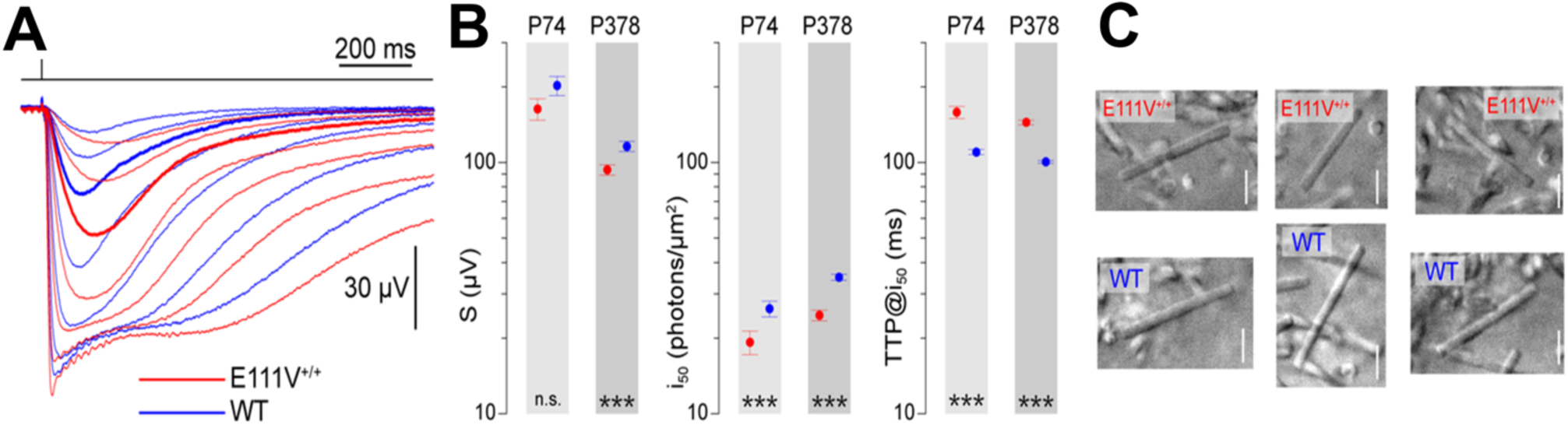
E111V^+/+^ mutants have more light-sensitive and slower rods. (**A**) Representative ex vivo ERG scotopic a-wave responses to flashes of increasing strength in ∼6 months old E111V^+/+^ (red) and WT retinas (blue) recorded at 37°C. Thicker traces highlight the differing responses to the same sub-saturating flash of the two animals, the mutant being more sensitive and slower. Flash strengths (photons/µm^2^): 2.89, 8.27, 18.9, 50.5, 151, 510, 1660. (**B**) Comparison of key flash response parameters in age-matched E111V^+/+^ (red) and WT mice (blue): saturating amplitude (S), half saturating flash strength (i_50_) and time to-peak at i_50_ (TTP@i_50_). Light gray: P74 on average; dark gray: P378 on average. Data are mean and SE. n.s.: not significant; *p < 0.05, **p < 0.01, ***p < 0.001. (**C**) Examples of rod outer segments (OSs) at P450 mutant and wild type mice, showing an apparently normal gross morphology. Live tissue imaged with a 63x water immersion objective, focusing on mechanically detached OSs for better imaging. Scale bars: 5 µm.

To determine whether this late functional decline was accompanied by overt structural deterioration of rod outer segments, we examined retinal ultrastructure at P450 (**Fig. 2C**). Rod outer segments appeared qualitatively indistinguishable between genotypes, displaying comparable length and morphology. These findings demonstrate that the E111V mutation produces persistent abnormalities in rod phototransduction long before extensive structural degeneration becomes apparent, identifying altered response dynamics as one of the earliest manifestations of severe GUCA1A-associated disease.

### E111V mice exhibit selective deficits in visually guided behavior

To determine whether the early photoreceptor dysfunction identified by ex vivo ERG translated into impaired visual performance at the organismal level, we assessed E111V⁺/⁺ mice using a battery of behavioral tests at P270, when retinal architecture remained largely preserved despite only modest ONL thinning and preserved lamination. The test battery included complementary assays probing visually guided depth perception, including the recently established Pole Descent Visual Cliff Task (*26*), together with tests evaluating locomotor activity, cognition, anxiety-like behavior and social interaction.

Behavioral abnormalities were remarkably selective. Whereas E111V⁺/⁺ mice performed comparably to WT controls in the Novel Object Recognition (NOR), Y-maze, Elevated Plus Maze (EPM) and Three-Chamber tests (**Supplementary Fig. S3**), indicating preserved cognitive function, anxiety-related behavior and social interaction, they consistently exhibited impaired performance in tasks requiring visually guided depth perception. In the classic Visual Cliff test, WT animals preferentially remained on the shallow (“safe”) side, whereas E111V⁺/⁺ mice spent significantly more time over the cliff and correspondingly less time in the shallow compartment (**Fig. 3B, C**), indicating impaired depth perception. This phenotype was independently confirmed by the Pole Descent Visual Cliff Task, in which mutant animals displayed significantly reduced landing accuracy on the shallow quadrant, as reflected by a lower percentage of correct trials (**Fig. 3E, F**; **Supplementary Movies 1** and **2**).

**Figure 3.**
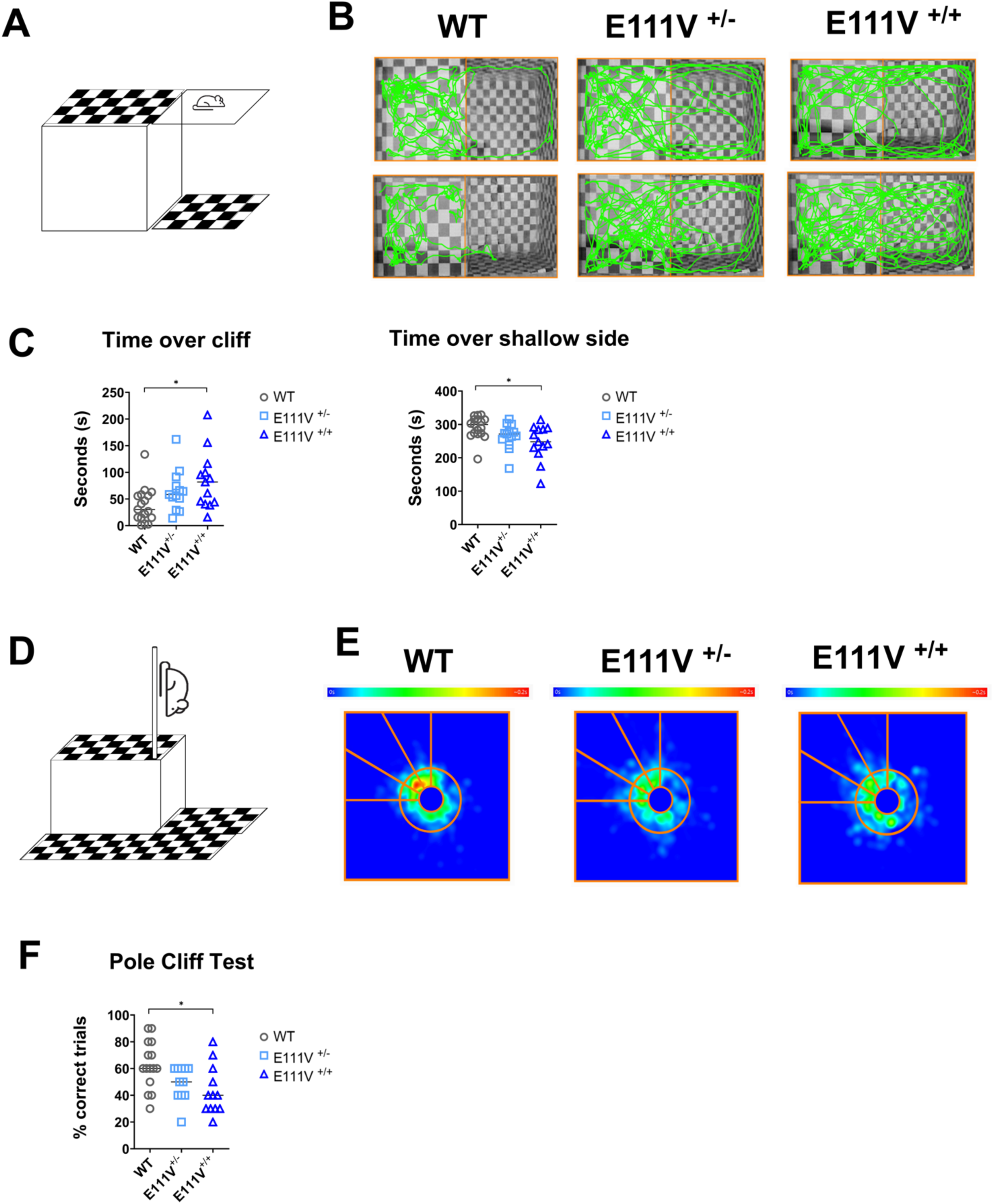
Depth perception is impaired in E111V^+/+^ mice. A–C. Visual Cliff test. (**A**) Schematic representation of the Visual Cliff test, (**B**) representative track plots of one animal per genotype (WT, E111V^⁺/⁻^, E111V^⁺/⁺^), (**C**) quantification of time spent over cliff and the shallow side. WT n = 17, E111V^⁺/⁻^ n = 13, E111V^⁺/⁺^ n = 13. **D–F** Pole Descent Visual Cliff task. (**D**) Schematic representation of Pole Descent Visual Cliff task, (**E**) the representative heatmaps of descent trajectories for each genotype. Heatmaps representing the position of the mouse data across all descent trials performed by WT ,E111V^+/−^ and E111V^+/+^ mice, with colour coding indicating cumulative time spent at each location (0 to ∼0.2 s; blue to red). The central circle marks the base of the pole, and the radial lines delimit the ’safe’ shallow zone where mice are expected to land; the remaining area constitutes the ’unsafe’ cliff zone. Warmer colours denote areas of greater cumulative occupancy, reflecting the preferred landing and dwelling location for each genotype. n = 10 trials for genotype. (**F**) quantification of correct choices for each genotype. WT n = 15, E111V^⁺/⁻^ n = 11, E111V^⁺/⁺^ n = 12. Statistical analysis: one-way ANOVA with Tukey’s multiple comparison test. *p < 0.05, **p < 0.01, ***p < 0.001.

E111V⁺/⁺ mice also displayed altered exploratory behavior in the Open Field Test (OFT) (**Supplementary Fig. S2** and **Supplementary Fig. S3B**), travelling longer distances at higher mean and maximum speeds while spending less time in the center of the arena. Although this pattern is commonly interpreted as anxiety-related thigmotaxis, the absence of differences in the EPM argues against an anxiety phenotype and instead suggests a vision-dependent compensatory locomotor strategy.

Together, these findings demonstrate that severe *GUCA1A*-associated disease selectively impairs visually guided behavior while preserving broader cognitive, emotional and social functions, indicating that the consequences of photoreceptor dysfunction extend beyond the outer retina.

### Retinal dysfunction propagates along the visual pathway in E111V mice

The selective behavioral deficits observed in E111V mice prompted us to ask whether visual information processing was impaired along the central visual pathway. We therefore recorded visually evoked potentials (VEPs) as local field potentials from the retinorecipient layers of the superior colliculus (SC) and the primary visual cortex (V1) in vivo at P120 and P270 (**Fig. 4**). WT recordings from WT animals (approximately 3 months old) were pooled and used as a common reference group. Visual responses in V1 progressively deteriorated with age in mutant animals. VEP amplitudes were significantly reduced in E111V⁺/⁺ mice at both P120 (0.82 ± 0.10 µV; P = 0.047) and P270 (0.54 ± 0.04 µV; P = 0.004) compared with WT controls (1.42 ± 0.29 µV), with a further significant decline between the two mutant age groups (P = 0.015) (**Fig. 4A, B**). Response latency showed a tendency towards prolongation at P270 but did not reach statistical significance either relative to WT (P = 0.065) or between mutant age groups (P = 0.068) (**Fig. 4C**).

**Fig. 4.**
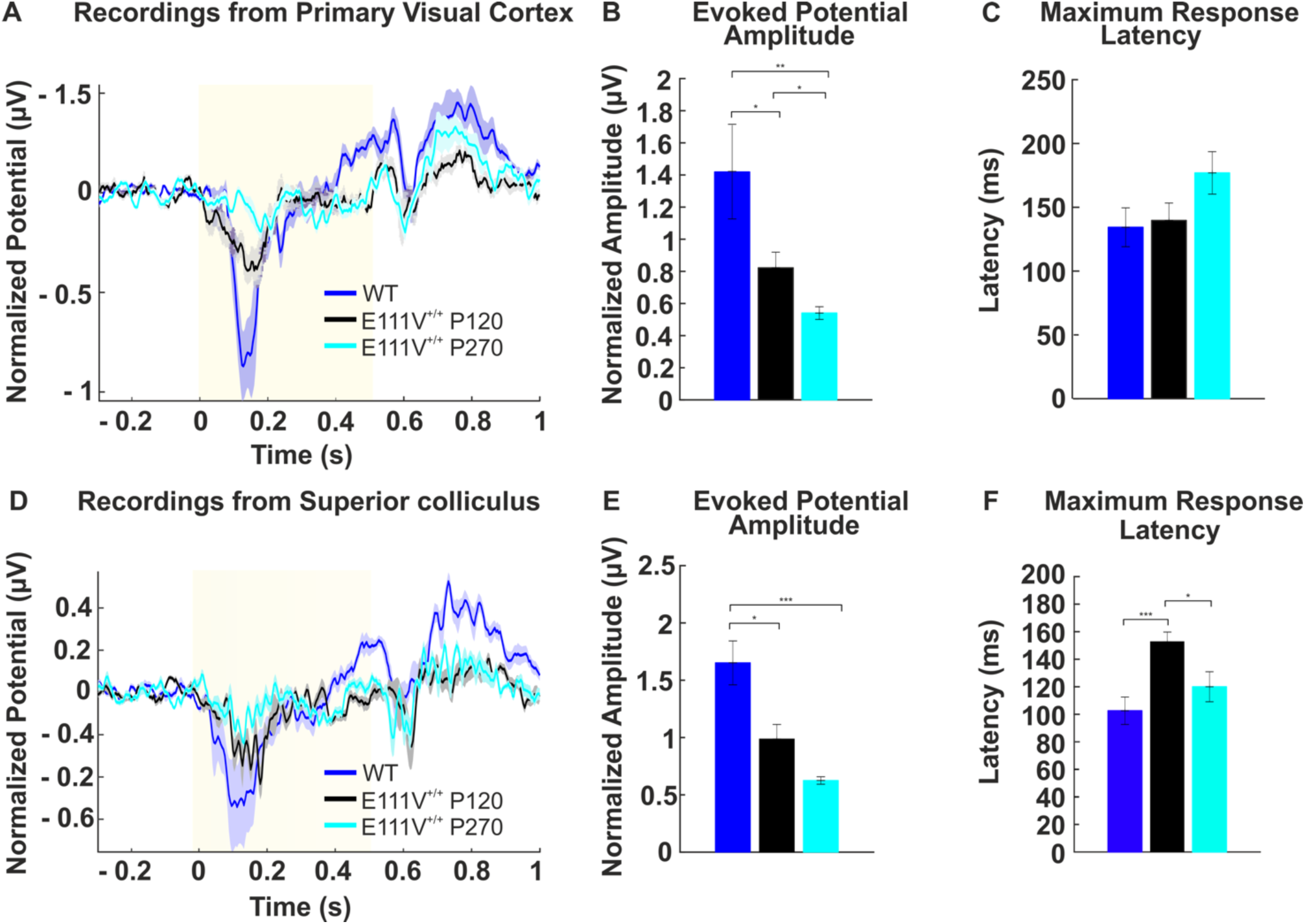
*In vivo* visual evoked potential (VEP) assessment in WT and E111V^+/+^ mice. Population VEP recordings from the primary visual cortex (V1) and superior colliculus (SC). (**A**) Representative population VEP traces recorded in V1 from WT (dark blue), E111V^+/+^ at P120 (black) and E111V^+/+^ at P270 (light blue) mice. (**B**) Mean VEP amplitudes in V1 across groups. (**C**) VEP response latency in V1 (no significant differences). (**D**) Representative population VEP traces recorded in SC from WT (dark blue), E111V^+/+^ at P120 (black) and E111V^+/+^ at P270 (light blue) mice. (**E**) Mean VEP amplitudes in SC across groups. (**F**) VEP response latency in SC; E111V^+/+^ mice show significantly prolonged latency compared with WT. Data are presented as mean ± SEM. Statistical comparisons by two-tailed Mann–Whitney U-test; *p < 0.05.

A similar reduction in response amplitude was observed in the superior colliculus (**Fig. 4D, E**). VEP amplitudes were significantly decreased in E111V⁺/⁺ mice at both P120 (0.99 ± 0.13 µV; P = 0.012) and P270 (0.63 ± 0.03 µV; P < 0.001) relative to WT animals (1.65 ± 0.19 µV), with a trend towards further decline with age that did not reach statistical significance (P = 0.055). In contrast to cortical responses, SC response latency was already significantly prolonged at P120 (152.8 ± 6.9 ms vs 102.6 ± 10.0 ms in WT; P < 0.001) and remained elevated at P270 (120.0 ± 10.9 ms; P = 0.033 vs WT; P = 0.430 vs P120) (**Fig. 4F**).

Together, these in vivo recordings demonstrate that functional abnormalities originating in photoreceptors propagate through the visual pathway, affecting both subcortical and cortical processing of visual information. The prolonged response latencies observed in the superior colliculus closely parallel the delayed rod photoresponse kinetics measured ex vivo, supporting the hypothesis that altered phototransduction contributes to impaired temporal processing throughout the visual system.

### E111V-GCAP1 disrupts the retinal distribution of key phototransduction signaling proteins

The propagation of functional abnormalities from photoreceptors to the retinal network prompted us to investigate whether the E111V mutation was accompanied by broader molecular reorganization within the retina. We therefore examined the retinal distribution of GCAP1 and three functionally related proteins involved in phototransduction signaling—GCAP2, the complementary guanylate cyclase-activating protein in murine retina (*27*); GC1, the principal effector enzyme responsible for cGMP synthesis (*4*); and retinal degeneration protein 3 (RD3), a key component of the GCAP1–GC1 regulatory complex (*28–31*). Immunohistochemical analyses were performed at P30 and P270 to compare early and later stages of disease progression.

In WT retinas at P30, GCAP1 localized predominantly to photoreceptors, displaying diffuse immunoreactivity throughout the outer segments (OS), stronger staining within the inner segments (IS), and a punctate signal in the outer plexiform layer (OPL) (**Fig. 5A**). E111V⁺/⁺ retinas exhibited a broadly similar distribution, although signal intensity was reduced within photoreceptor layers and the OPL, with additional weak immunoreactivity detectable in the inner plexiform layer (IPL) (**Fig. 5B**). At P270, GCAP1 staining decreased in both genotypes; however, mutant retinas displayed a striking redistribution toward the ganglion cell layer (GCL), where immunoreactivity became markedly enriched (**Fig. 5C, D**).

**Fig. 5.**
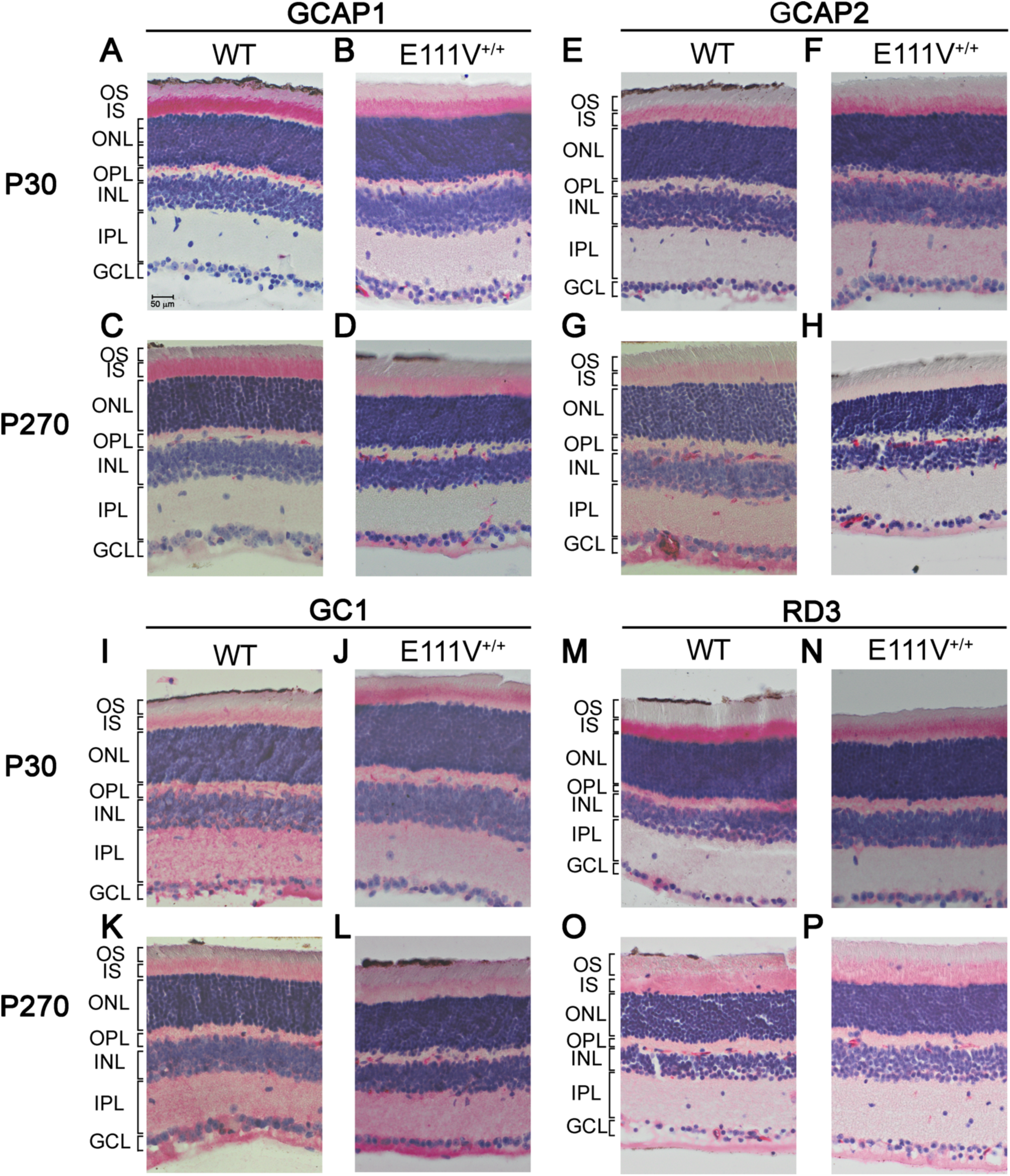
Retinal immunolocalization of GCAP1, GCAP2, GC1, and RD3 in WT and E111V^+/+^ mice. Representative images from n = 6 mice per genotype at P30 and P270 for GCAP1 (**A–D**), GCAP2 (**E–H**), GC1 (**I–L**) and RD3 (**M–P**). Images were acquired with an Evident APEX microscope (20× objective). Scale bar: 100 µm.

GCAP2 displayed a distinct but partially overlapping localization pattern. In WT retinas at P30, immunoreactivity was concentrated within photoreceptor IS, with weaker staining in the OS and minimal labeling of the GCL (**Fig. 5E**). In E111V⁺/⁺ retinas, GCAP2 distribution closely resembled that of GCAP1 (**Fig. 5F**), suggesting an early adaptive response of the complementary GCAP isoform. By P270, GCAP2 immunoreactivity became weak throughout WT retinas while remaining relatively enriched in the GCL (**Fig. 5G**). In contrast, E111V⁺/⁺ retinas exhibited a marked reduction of GCAP2 labeling across all retinal layers except the GCL, where immunoreactivity persisted (**Fig. 5H**).

GC1 exhibited a broader retinal distribution than GCAP1. At P30, WT retinas showed a gradient of immunoreactivity extending from the outer to the inner retina, with prominent labeling in the IS, OPL, IPL and GCL (**Fig. 5I**). In contrast, E111V⁺/⁺ retinas displayed an inverted distribution pattern characterized by stronger immunoreactivity within the photoreceptor layers and reduced labeling toward the GCL (**Fig. 5J**). Punctate GC1 staining was also evident in the OPL, closely resembling the distribution observed for GCAP1. By P270, GC1 became more uniformly distributed throughout WT retinas (**Fig. 5K**), whereas E111V⁺/⁺ retinas exhibited a progressive redistribution toward the inner retina, with particularly intense immunoreactivity within the GCL (**Fig. 5L**).

RD3 showed a distribution largely overlapping that of GCAP1 and GC1 at early stages. In WT retinas at P30, RD3 immunoreactivity was concentrated within photoreceptor IS and the OPL, where it displayed a characteristic punctate pattern, with only weak labeling of the GCL (**Fig. 5M**). E111V⁺/⁺ retinas exhibited a similar overall distribution but with reduced signal intensity within the IS (**Fig. 5N**). By P270, RD3 became more uniformly distributed throughout photoreceptor layers, OPL, IPL and GCL in both genotypes, with no obvious mutation-dependent redistribution.

Collectively, these observations indicate that the E111V mutation reshapes the retinal distribution of multiple components of the GCAP1/GC1 signaling machinery, extending well beyond the mutant protein itself. The coordinated redistribution of GCAP1, GCAP2 and GC1 across retinal layers is consistent with widespread molecular reorganization accompanying the emergence of retinal network dysfunction. Although the intense GCL immunoreactivity observed at P270 could partly reflect age-related accumulation of macromolecular material or lipofuscin, these findings support the view that severe GUCA1A-associated disease involves progressive alterations extending beyond photoreceptor signaling alone.

### Transcriptomic profiling reveals progressive disruption of retinal homeostasis in E111V mice

To identify molecular processes associated with the retinal network dysfunction observed in E111V⁺/⁺ mice, we performed transcriptomic profiling of whole eyecup preparations, encompassing the neural retina and closely apposed retinal pigment epithelium (RPE), from WT and E111V⁺/⁺ animals at P30 and P270. Sequencing yielded an average of 38 million reads per sample, with over 95% passing quality control (**Supplementary Data 2**). Principal component analysis revealed clear genotype- and age-dependent transcriptional separation across conditions (**Supplementary Fig. S4**). Differential expression analysis identified 177 differentially expressed genes (DEGs) at P30, including 45 upregulated and 132 downregulated transcripts relative to WT (**Supplementary Figs. S4, S5** and **Supplementary Data 3**). By P270, transcriptional dysregulation had markedly expanded to 1148 DEGs, including 451 upregulated and 697 downregulated transcripts (**Supplementary Figs. S6, S7** and **Supplementary Data 4**). Longitudinal analyses further revealed age-dependent expression changes in both genotypes, with distinct transcriptional trajectories in E111V⁺/⁺ and WT retinas (**Supplementary Data 5, 6**).

To define the biological processes underlying these transcriptional changes, we performed gene ontology (GO) enrichment analysis and Gene Set Enrichment Analysis (GSEA). Enriched terms revealed genotype- and age-specific perturbations involving visual perception, synaptic translation and metabolism. Notably, the E111V⁺/⁺ P30-to-P270 comparison showed enrichment of terms and gene sets associated with visual perception, whereas the corresponding WT comparison was enriched for mitochondrial-related pathways, indicating divergent aging-related transcriptional responses between genotypes (**Supplementary Figs. S8–S11** and **Supplementary Data 7,8**). We next examined genes involved in retinal function and development, synaptic signaling, metabolism, inflammation, cell death and cytoskeletal organization to identify molecular signatures associated with disease progression (**Fig. 6**). Among genes involved in retinal homeostasis, *Mertk* emerged as one of the earliest and most persistent transcriptional alterations (**Fig. 6B**). *Mertk*, which is predominantly expressed in the RPE, mediates phagocytic clearance of photoreceptor outer segments and is essential for retinal homeostasis (*32, 33*). In E111V⁺/⁺ mice, *Mertk* was already significantly downregulated at P30 (log2FC = –1.17; adjusted P = 2.16 × 10⁻⁷), and this reduction persisted at P270 (log2FC = –1.14; adjusted P = 4.73 × 10⁻⁵) (**Fig. 6B**). Although bulk transcriptomics cannot distinguish whether this early suppression reflects intrinsic RPE dysfunction or a secondary response to altered photoreceptor outer-segment turnover, its early and sustained nature implicates impaired clearance mechanisms among the earliest molecular alterations associated with severe *GUCA1A* disease.

**Fig. 6.**
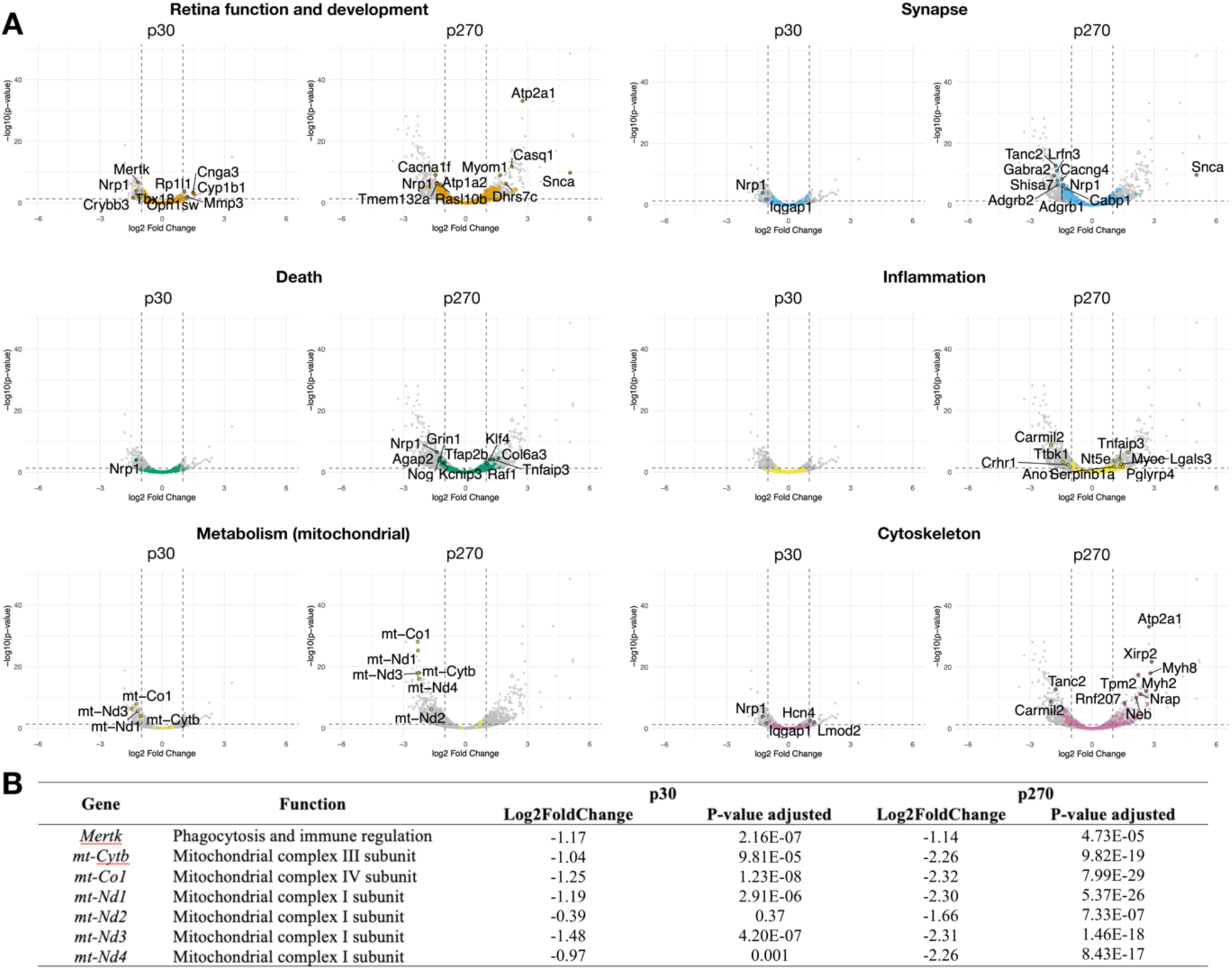
RNAseq analysis results. (**A)** Volcano plots of differential gene expression for the inter-genotype comparisons between E111V^+/+^ and WT mice at P30 and P270. For each comparison, genes belonging to selected functional categories are highlighted in color, while all other genes are shown in grey. Metabolism (mitochondrial) refers to genes involved in the respiratory electron transport chain. The plots display log2 fold change versus −log10 adjusted p-value. (**B**) Differential gene expression analysis between E111V^+/+^ and WT samples at timepoint P30 and P270. For each selected gene, the table reports the log2 fold change between E111V^+/+^ and WT and adjusted p-value.

Because persistent *Mertk* downregulation suggested impaired photoreceptor outer-segment clearance, we next examined retinal ultrastructure by transmission electron microscopy (TEM) to determine whether these early molecular alterations were accompanied by structural abnormalities of photoreceptors (**Fig. 7**). WT retinas displayed the expected orderly lamellar organization of photoreceptor outer segments. In contrast, E111V⁺/⁺ retinas exhibited marked ultrastructural abnormalities, including densely packed and disorganized outer-segment discs together with frequent accumulation of electron-dense vesicular structures within the inner segments. Although the molecular identity of these vesicles will require future immunogold characterization, their presence is consistent with impaired outer-segment turnover. Importantly, these profound ultrastructural abnormalities were observed despite the relatively preserved retinal architecture detected by light microscopy (**Fig. 1** and **Supplementary Fig. S1**), indicating that conventional histology substantially underestimates the extent of early photoreceptor pathology. Together, these ultrastructural findings provide morphological support for the early *Mertk* downregulation identified by transcriptomic profiling and are consistent with impaired RPE-mediated photoreceptor maintenance during the earliest stages of disease progression.

**Fig. 7.**
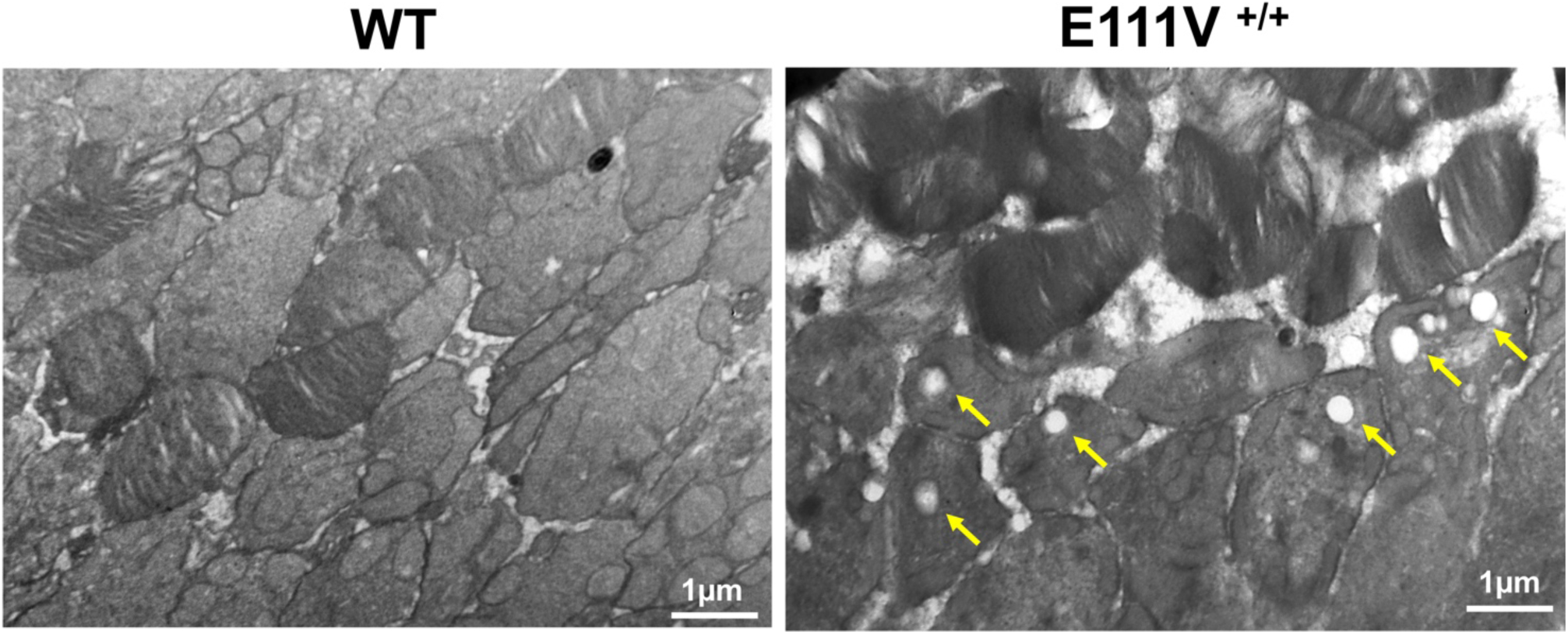
Transmission electron microscopy (TEM). Two representative images of WT (**A**) and E111V^+/+^ (**B**) photoreceptors at P270. The yellow arrows indicate vesicle-like structures. Scale bar: 1µm

Beyond the early alteration of *Mertk*, transcriptomic profiling revealed a broader and progressive molecular reprogramming of the mutant retina. At P30, transcriptional alterations remained relatively restricted and primarily affected genes associated with retinal function, synaptic signaling and mitochondrial metabolism, whereas inflammatory and cell-death pathways showed minimal activation (**Fig. 6**). By P270, this molecular signature had evolved into widespread dysregulation encompassing retinal signaling, synaptic organization, mitochondrial metabolism, inflammation, cell death and cytoskeletal organization, indicating progressive disruption of retinal homeostasis. Synaptic pathways showed progressive dysregulation of genes involved in synaptic organization and neurotransmission, whereas cytoskeletal remodeling also became increasingly prominent at the later disease stage (**Fig. 6B**).

Mitochondrial dysfunction emerged as one of the earliest and most persistent molecular signatures. Already at P30, multiple mitochondrial-encoded genes involved in oxidative phosphorylation, including *mt-Co1*, *mt-Nd1*, *mt-Nd3* and *mt-Cytb*, were significantly downregulated (**Fig. 6B**). This signature became markedly more pronounced at P270, when mitochondrial respiratory chain genes exhibited dramatic suppression (for example, *mt-Cytb*, log2FC = –2.26; *mt-Nd1*, log2FC = –2.30; *mt-Nd3*, log2FC = –2.31), together with additional downregulation of *mt-Nd2* and *mt-Nd4*, consistent with progressive impairment of mitochondrial respiratory function. In contrast, inflammatory and cell-death pathways showed only limited activation at P30 but became prominent at P270, indicating that neuroinflammatory responses represent a relatively late component of disease progression rather than an initiating event.

### Wild-type GCAP1 partially restores rod photoresponse dynamics in E111V mice

Because the functional abnormalities identified in E111V photoreceptors arise from altered GCAP1 regulation of retinal guanylate cyclase, we next asked whether supplementation with recombinant wild-type (WT) GCAP1 could partially restore rod photoresponse dynamics. Previous work from our laboratories demonstrated that recombinant E111V-GCAP1 readily enters WT photoreceptors following ex vivo incubation and slows their photoresponse kinetics (*24*).

As a control experiment, recombinant E111V-GCAP1 was delivered to E111V⁺/⁺ retinas. One retina from each animal was incubated with 3.2 µM recombinant protein and the contralateral retina with Ames’ medium alone. No systematic effects on either light sensitivity or response kinetics were observed (**Fig. 8A**; n = 8 animals), indicating that the endogenous mutant protein already dominates rod photoresponse regulation under these conditions. We next examined whether supplementation with recombinant WT-GCAP1 could partially reverse the electrophysiological phenotype. One retina from each E111V⁺/⁺ mouse was incubated with 3.2 µM WT-GCAP1, whereas the contralateral retina received vehicle alone. WT-GCAP1 consistently shifted rod response parameters toward WT values, producing a reproducible partial correction of the mutant electrophysiological phenotype (**Fig. 8B**; n = 25 animals). These findings provide proof-of-concept that the early functional abnormalities associated with severe *GUCA1A* disease remain amenable to biochemical modulation, supporting the existence of a therapeutic window before irreversible structural degeneration.

**Fig. 8.**
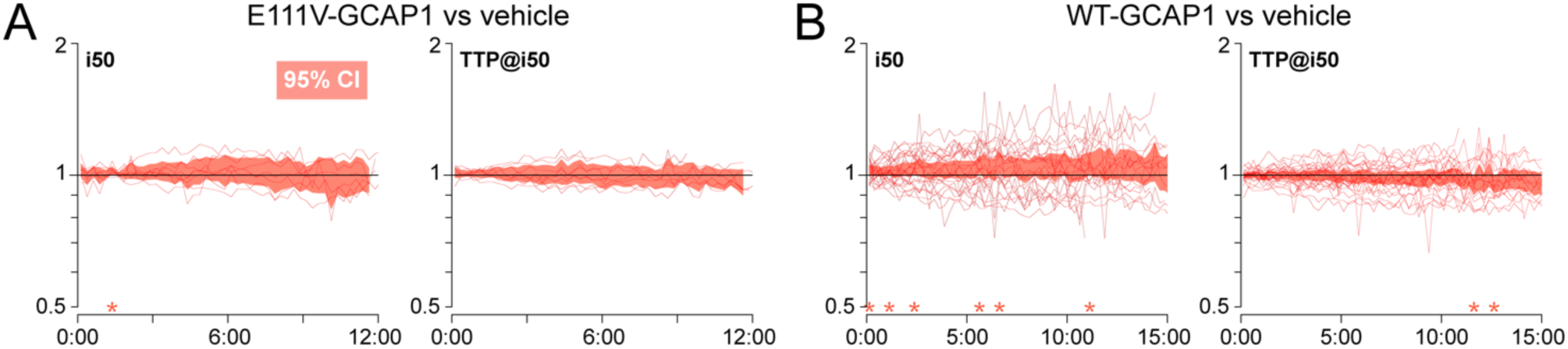
Effects of mutant and wild type recombinant GCAP1 on the scotopic flash responses of E111V^+/+^ mice. (**A**) Time course of the relative effect on rod i_50_ and TTP@i_50_ (see legend of fig. 2) of incubating the retinas of E111V^+/+^ mutants with 3.2 µM E111V-GCAP1. Each line represents the raw data from one animal, while the filled area is the 95% confidence interval (CI); temperature 37°C. (**B**) Effects of incubating E111V^+/+^ mutant retinae with 3.2 µM WT-GCAP1. The trend toward values greater than unity of i_50_ would imply a decrease in light sensitivity, while the opposite trend shown by TTP@i_50_ suggests a quickening of response kinetics. *: p < 0.05. All recordings were made at 37°C.

## DISCUSSION

The relationship between photoreceptor dysfunction and progressive vision loss remains incompletely understood in inherited retinal degeneration. Using a physiologically faithful knock-in mouse carrying the clinically most severe GCAP1 variant, we show that severe *GUCA1A*-associated cone-rod dystrophy is characterized by profound functional abnormalities that emerge while retinal architecture remains largely preserved. This dissociation between function and structure extends from altered photoreceptor response dynamics to visually guided behavior and central visual processing, identifying retinal network dysfunction as an early feature of disease progression rather than a secondary consequence of photoreceptor degeneration.

Contrary to expectations, the marked biochemical severity of the E111V mutation did not translate into rapid structural degeneration. ONL thickness remained essentially preserved at P30 and P120, with significant thinning apparent only at P270 in homozygous animals (**Fig. 1**). This slow progression resembles that of the E155G (*21*) and L151F (*22*) *GUCA1A* mouse models and contrasts sharply with rapidly degenerating models such as P23H rhodopsin transgenic mice (*34*) and the rd10 mouse (*35*), suggesting that species-specific compensatory mechanisms buffer photoreceptor survival in mice. Consistent with this, scotopic ERG responses — virtually absent in E111V-GCAP1 patients by the third decade of life (*16*) — remained readily detectable in age-matched E111V⁺/⁺ mice despite markedly increased light sensitivity and slowed photoresponse kinetics (**Fig. 2**). This combination is best explained by weakened Ca²⁺-dependent negative feedback on retinal guanylate cyclase rather than by elevated dark cGMP alone: because the fractional rod photoresponse is governed mainly by the dynamics of Ca²⁺-dependent cyclase regulation and is relatively insensitive to the absolute dark cGMP concentration, the rightward shift in Ca²⁺ sensitivity of E111V-GCAP1 prolongs response integration time, producing parallel increases in flash sensitivity and time-to-peak. This is consistent with our kinetic modelling of E111V (*25, 36*) and with recordings from the Y99C-GCAP1 mouse, in which constitutive cyclase activation similarly produced larger and slower single-photon responses while preserving overall rod function (*20*). The persistence of dynamically regulated rod responses — despite the absence of cone recordings, a limitation of this study — indicates that residual Ca²⁺-dependent cyclase regulation is maintained, likely supported by mouse GCAP2, which unlike its human orthologue retains the ability to regulate GC1 (*27, 37*). This species-specific difference probably explains why clinically severe heterozygous *GUCA1A* mutations require homozygosity to produce a robust phenotype in mice. Rather than a limitation, the slower progression offers a unique opportunity to investigate the earliest pathogenic events preceding overt photoreceptor loss — a stage largely inaccessible in affected patients.

The preservation of retinal structure contrasts strikingly with the magnitude of the functional phenotype across the visual system. Behaviorally, E111V⁺/⁺ mice performed normally in tests of recognition memory, spatial working memory, anxiety-like behavior and sociability, but were consistently impaired in tasks requiring visually guided depth perception (**Fig. 3** and Supplementary **Figs. S2, S3**). This selectivity argues against generalized neurological dysfunction and points to specific disruption of visual processing. The deficit was paralleled by progressive reductions in visually evoked responses in both the superior colliculus and primary visual cortex (**Fig. 4**), showing that the consequences of altered photoreceptor signaling propagate beyond the outer retina. Notably, the age-dependent decline in VEP amplitude and the early prolongation of superior colliculus response latency closely mirror the delayed rod photoresponse kinetics measured ex vivo, suggesting that abnormal phototransduction may actively shape the timing of visual information processing along the pathway rather than being merely associated with downstream dysfunction.

These observations broaden the prevailing view of *GUCA1A*-associated disease. Rather than a disorder confined to phototransduction within photoreceptors, defective GCAP1 signaling has functional consequences that reach retinal circuits and central visual processing well before extensive structural degeneration becomes apparent. Whether this reflects trans-synaptic propagation of altered photoreceptor output, adaptive changes within inner-retinal circuitry, or a combination of both remains to be established. Our immunohistochemistry supports this broader interpretation. Although the overall distribution of GCAP1, GCAP2, GC1 and RD3 was largely preserved at P30, mutant retinas showed progressive age-dependent redistribution of GCAP1 and GC1 toward the ganglion cell layer (**Fig. 5**). We recently showed that the GCAP1–RD3 interaction is strongly Ca²⁺-dependent — Ca²⁺-bound GCAP1 binds RD3 with micromolar affinity, whereas the Mg²⁺-bound form interacts far more weakly — and that E111V uniquely abolishes this interaction, unlike other disease-associated variants such as D100G, N104H and E155G, which retain RD3 binding with altered kinetics (*31*). Because GCAP1, GC1 and RD3 normally co-localize in photoreceptor inner segments and synaptic terminals, where local Ca²⁺ favors complex formation, disruption of this regulatory node is likely to perturb GC1 trafficking and local signaling homeostasis. Together with the absence of GCAP2-mediated GC1 regulation in the human retina (*37*), this framework may explain why the pathological consequences of E111V are considerably more severe in patients than in mice.

Convergent transcriptomic and ultrastructural evidence indicates disruption of retinal homeostasis from the earliest disease stages. Rather than isolated molecular alterations, transcriptomic profiling revealed a coordinated progression from early defects in retinal maintenance and mitochondrial metabolism to broader dysregulation of synaptic organization, inflammation and cytoskeletal pathways (**Fig. 6**), closely paralleling the physiological progression across retinal and central visual function. Among the earliest and most persistent alterations, *Mertk* — which encodes the receptor tyrosine kinase required for RPE-mediated phagocytosis of shed photoreceptor outer segments (*32, 33*) — was a particularly compelling candidate: loss-of-function *MERTK* mutations cause severe early-onset retinitis pigmentosa in humans and rapid photoreceptor degeneration in experimental models (*32, 33*). Its sustained downregulation from P30 onward suggests that impairment of the RPE–photoreceptor axis is among the earliest consequences of severe GCAP1 dysfunction, although bulk transcriptomics cannot distinguish primary RPE dysregulation from secondary responses to altered photoreceptor turnover. Transmission electron microscopy provided independent structural support: despite the relatively preserved architecture seen by conventional histology, E111V⁺/⁺ retinas displayed profound ultrastructural abnormalities, including disorganized outer-segment disc membranes and electron-dense vesicular structures within photoreceptor inner segments. These are consistent with defective outer-segment renewal and impaired clearance — whose molecular identity remains to be established — and indicate that conventional light microscopy substantially underestimates the extent of early photoreceptor pathology.

Mitochondrial dysfunction was another prominent molecular signature: downregulation of multiple mitochondrially encoded respiratory-chain genes was already detectable at P30 and became markedly more pronounced with age, consistent with progressive impairment of oxidative phosphorylation. Because photoreceptors have among the highest metabolic demands of any neuronal cell type, this bioenergetic decline is likely to amplify the consequences of defective phototransduction by reducing the capacity to maintain ionic homeostasis, synaptic transmission and outer-segment renewal. In contrast, inflammatory and cell-death pathways became prominent only at later stages, suggesting that neuroinflammation is more likely a secondary consequence of chronic retinal dysfunction than an initiating event.

Finally, the partial restoration of rod photoresponse dynamics by recombinant wild-type GCAP1 provides proof-of-concept that the earliest functional abnormalities associated with severe *GUCA1A* disease remain biologically modifiable. Performed under acute ex vivo conditions and not intended to demonstrate therapeutic efficacy, these experiments indicate that the mutant electrophysiological phenotype is not irreversibly fixed during early disease but reflects a dynamic stoichiometric imbalance susceptible to competition by wild-type GCAP1 (**Fig. 8**). This is consistent with our previous observations that recombinant GCAP1 efficiently penetrates rod photoreceptors and retinal ganglion cells in murine and porcine retinal explants — with intracellular retention enhanced by the retina-specific myristoylated form — and that liposomal formulations engineered to mimic outer-segment membranes provide controlled, sustained release of recombinant protein (*24, 38*), supporting the feasibility of protein-based delivery strategies for retinal disease. This rescue is unlikely to reflect simple competition between endogenous and exogenous GCAP1 alone. Because E111V abolishes the GCAP1–RD3 interaction, as noted above (*31*), delivery of recombinant wild-type GCAP1 may not only rebalance GC1 regulation but also restore this critical regulatory complex, providing a plausible mechanism for the observed improvement in rod photoresponse dynamics. As recombinant wild-type GCAP1 competes with mutant protein independently of the specific amino acid substitution, the strategy could in principle apply across the spectrum of dominant *GUCA1A*-associated cone-rod dystrophies rather than being restricted to E111V. Whether comparable — or even greater — functional correction can be achieved in less severe variants, or following chronic in vivo delivery, remains an important question for future investigation.

Collectively, our findings support a revised view of severe *GUCA1A*-associated cone-rod dystrophy as a progressive disorder of retinal network function rather than a disease confined to phototransduction. By demonstrating that functional impairment precedes overt structural degeneration, identifying disruption of retinal homeostasis as a plausible driver of disease progression, and providing evidence that early functional abnormalities remain biologically modifiable, this study defines the earliest stages of disease and highlights an early window for therapeutic intervention before irreversible retinal degeneration becomes established.

## MATERIALS AND METHODS

### Animals

All animal procedures were approved by the University of Verona’s Animal Care and Use Committee (CIRSAL) and the Italian Ministry of Health, and by the 2nd Local Ethical Committee in Warsaw (permission no. WAW2/009/2025), in compliance with the European Communities Council Directive 2010/63/EU, NIH guidelines, and the ARVO Statement for the Use of Animals in Ophthalmic and Vision Research. Animals were maintained under a 12 h light/dark cycle with controlled temperature and humidity and ad libitum access to food and water.

### Generation of knock-in mice

Knock-in (KI) mice carrying the *GUCA1A* p.E111V variant (c.332A>T;333G>T; GAG>GTT) and a silent mutation p.L113= (CTC>TTA) were generated by Cyagen Biosciences (Santa Clara, CA, USA) via CRISPR/Cas9 co-injection of targeting gRNAs and a donor oligonucleotide into fertilized C57BL/6J oocytes. Founder mice, obtained homozygous, were crossed with WT C57BL/6J mice (The Jackson Laboratory, Bar Harbor, ME, USA) to establish a heterozygous colony. WT, E111V^+/−^ and E111V^+/+^ offspring were genotyped by PCR and analyzed at postnatal days 30–270 (P30–P270).

### Retinal morphology

For retinal morphology analyses, six animals per genotype (WT, E111V^+/−^ and E111V^+/+^) were collected at P30–P270. Mice were anesthetized with isoflurane and euthanized by cervical dislocation. Eyes were enucleated, the anterior segment removed along the *ora serrata*, and the resulting eyecups fixed in 10% neutral buffered formalin (NBF) for 1 h at room temperature (RT). After PBS washes, eyecups were cryoprotected through sequential sucrose gradients (10%, 20%, 30%), equilibrated in a 1:1 OCT/30% sucrose mixture, snap-frozen in liquid nitrogen, and stored at −80°C. Sagittal cryosections (10–14 µm) were cut on a cryostat, counterstained with DAPI (Invitrogen, 1:1000), and mounted with Dako fluorescent mounting medium (Agilent Technologies, cat. no. S3023). Images were acquired on an Olympus microscope with a 20× objective. Outer nuclear layer (ONL) thickness was measured in ImageJ across six fields per section centered on the optic nerve and averaged per animal; values were normalized to WT as fold change.

### Immunohistochemistry

Eyes from WT and E111V^+/+^ mice were enucleated, punctured, and fixed overnight in 10% NBF at 4°C. Corneas and lenses were removed and eyecups post-fixed for 2 h at RT, dehydrated through a graded ethanol series, cleared in xylene, and paraffin-embedded. Sections (7 µm) were mounted on poly-L-lysine-coated slides, deparaffinized in xylene, and rehydrated through graded ethanol. Antigen retrieval was performed in citrate buffer (pH 6.0, 85°C, 20 min). Sections were blocked in 20% normal goat serum (NGS) with 0.5% Tween-20 in PBS (1 h, RT) and incubated overnight at 4°C with primary antibodies against GC1 (Novus Biologicals, NBP3-12211), GCAP1 (Novus Biologicals, NBP3-17717), GCAP2 (Novus Biologicals, NBP2-68721), and RD3 (Invitrogen, PA5-83117), diluted in 0.5% BSA-PBS. Detection was performed on a Leica BOND-III automated system using a BOND-PRIME Polymer AP Detection System, followed by hematoxylin counterstaining. For morphological assessment, adjacent sections were stained with hematoxylin and eosin (H&E; Bio-Optica kit). Images were acquired on an Evident APX100-HCU microscope with a 20× objective.

### Immunofluorescence

Paraffin sections were processed as described above through the antigen retrieval step, then washed in PBS–0.5% Triton X-100 and blocked for 1 h at RT in PBS containing 20% donkey serum, 50 mM glycine, 5% BSA, 0.05% Tween-20, and 0.1% Triton X-100. Sections were incubated overnight at 4°C with anti-PKCα antibody (Sigma-Aldrich, P4334) and Alexa Fluor 488-conjugated PNA lectin (Molecular Probes, L-21409) in PBS–10 mM glycine–0.05% Tween-20–0.1% Triton X-100, followed by Alexa Fluor 647-conjugated secondary antibody (1 h, RT). Nuclei were counterstained with DAPI and slides were mounted with Dako fluorescent mounting medium. Images were acquired on an Evident APX100-HCU microscope with a 20× objective.

### RNA Extraction and 3′ mRNA Sequencing Library Preparation

Frozen eyecups for WT and homozygous mice (n=3 per group; contralateral eyecup for each of the samples used in immunohistochemistry experiments) were homogenized on ice by sonication (50% amplitude; 2-second pulses with 2-second intervals; total processing time: 10 minutes). Total RNA was extracted using the RNeasy Mini Kit (Qiagen, Cat. No. 74104) according to the manufacturer’s instructions. RNA concentration and purity were assessed spectrophotometrically using a NanoDrop One spectrophotometer (Thermo Fisher Scientific), and RNA integrity was determined using the DNF-471-33 - SS Total RNA kit on a Fragment Analyzer instrument (Agilent Technologies) prior to library preparation. Strand-specific 3′ mRNA sequencing libraries were constructed using the QuantSeq 3′ mRNA-Seq Library Prep Kit FWD (Lexogen) following the manufacturer’s protocol. Sequencing was performed on an Illumina NextSeq 2000 platform in single-read 75 bp mode (75SR), yielding an average of approximately 38 million reads per sample. Library preparation and sequencing were performed at the Centro Piattaforme Tecnologiche (University of Verona, Verona, Italy).

### Reads processing, alignment, and transcript quantification

Raw reads were quality-checked, aligned to the *Mus musculus* reference genome, and processed for transcript quantification. In detail, the quality of raw reads was assessed using FastQC v0.12.1(*39*), and Cutadapt v2.8 (*40*) was used to trim adapter sequences. For transcript quantification, reads were aligned to the *M. musculus* GRCm39 primary assembly reference genome downloaded from Gencode (downloaded on 17 October 2024 from https://ftp.ebi.ac.uk/pub/databases/gencode/Gencode_mouse/release_M36) using STAR v2.7.11b (*41*) Transcripts were quantified using the primary assembly basic annotation GTF file and the parameter --quantMode GeneCounts.

### Differential Expression, Gene Ontology, and Gene Set Enrichment Analyses

Differential expression analysis (DEA) and functional enrichment were performed to identify GUCA1A genotype- and age-dependent transcriptional changes in mice, highlighting perturbed biological pathways. DEA was performed using the DESeq2 package v1.46.0 (*42*) considering two sets of comparisons: (1) inter-genotype (i.e., E111V^+/+^ vs WT at P30 and at P270), and (2) intra-genotype longitudinal contrasts (i.e., P270 vs P30 in WT and in E111V^+/+^ mice). The WT genotype and the P30 timepoint were used as reference levels for the inter-genotype and intra-genotype analyses, respectively. Gene Ontology (GO) enrichment analysis and Gene Set Enrichment Analysis (GSEA) were carried out using the enrichGO and gseGO functions of the clusterProfiler R package v4.14.4 (*43*). GO and gene set annotations for *M. musculus* were retrieved using the org.Mm.eg.db package v3.20.0. For GO enrichment analysis, the list of significantly modulated genes identified by DEA was used as input; whereas GSEA was performed using a list of all expressed genes ranked by their log2FoldChange, considering gene sets with a minimum size of 3 and a maximum size of 800. In addition to the unbiased differential expression and enrichment analyses, targeted analyses focusing on specific biological processes of interest (retina function and development, synapses, cell death, inflammation, metabolism, and cytoskeleton) were performed. Gene sets were defined based on curated functional categories and keyword-based annotations relevant to the biological questions addressed in this study. In particular, due to the heterogeneity and numerosity of the genes constituting the metabolism process, further analysis was limited to a subset representing those involved in the respiratory electron transport chain (GO:0022904). These gene sets were then examined within the differential expression results to assess E111V^+/+^ genotype- and age-dependent transcriptional changes specifically affecting these processes. The full list of biological categories and associated keywords used to define the targeted gene sets is provided in **Supplementary Data 1**. All plots were generated with ggplot2 v3.5.1 (*44*), and all analyses were carried out in an R v4.4.1 environment.

### Recombinant protein production

Recombinant myristoylated WT-GCAP1 and E111V-GCAP1 were produced in *Escherichia coli* BL21(DE3) cells co-transformed with a pET-11a plasmid encoding the respective human GCAP1 variant and a pBB131 vector encoding *S. cerevisiae* N-myristoyltransferase (yNMT), required for co-translational N-terminal myristoylation, as previously described (*16, 31*), Briefly, myristic acid (50 µg/ml in 50% EtOH) was added to the growth medium at OD₆₀₀ = 0.4, and protein expression was induced with 1 mM IPTG at OD₆₀₀ = 0.6 for 4 h at 37°C. GCAP1 variants were extracted from inclusion bodies by denaturation in 6 M guanidine-HCl and refolded by dialysis against 20 mM Tris-HCl pH 7.5, 150 mM NaCl, 14 mM β-mercaptoethanol. Proteins were purified by Size Exclusion Chromatography (SEC; HiPrep 26/60 Sephacryl S-100 HR, GE Healthcare) followed by Anion Exchange Chromatography (AEC; HiPrep Q HP 16/10, GE Healthcare) using a linear 0–1 M NaCl gradient. Protein concentration was determined by Bradford assay(*45*) using a GCAP1-specific calibration curve, and purity assessed by 15% SDS-PAGE. Purified proteins were lyophilized and stored at −80°C until use. For details on the retinal delivery protocol, see the Electroretinography section and refs.(*24, 38, 46*).

### Electroretinography

To characterize photoreceptor function in the GCAP1 mutant, monitor disease progression, and test the effects of recombinant GCAP1 incubation, we performed ex vivo ERG recordings on isolated retinas. These were obtained using a custom-designed apparatus that permits simultaneous acquisition of the pharmacologically isolated scotopic a-wave responses (*47*). E111V^+/+^ and C57BL/6J mice of both sexes were dark-adapted for >1 hour and euthanized by intraperitoneal injection of ketamine (∼0.4 mg/g) + xylazine (∼0.06 mg/g). Retinae were isolated in incubation solution, made to adhere on filter paper (SMWP02500; Sigma-Merck) with their photoreceptors facing upwards, and placed in recording wells (2 mL incubation solution/retina). Transretinal potentials were measured during incubation at 37°C in a humidified 95% O_2_/5% CO_2_ atmosphere. Signals were filtered (0.02–100 Hz), amplified and acquired at 5 kHz with a Digidata 1322B and pClamp 9 software (Molecular Devices, San Jose, CA). Sequences of full-field flashes were delivered at 15 min intervals with a green LED (505 nm peak) attenuated using neutral density filters. Flash strength was adjusted by varying the LED drive current and/or duration. Each sequence consisted of the following flashes (strength at the retina [photons/µm^2^]|duration [ms]|repetitions [N]| interval [s]): 2.89|1|12|3, 8.27|1|10|3.6, 18.9|1|8|4.5, 50.5|1|6|6, 151|3|6|6, 510|6|4|9, 1660|15|3|18. The incubation solution was Ames’ medium (A1420; Sigma-Merck) supplemented with 40 µM AP4 (0101; Tocris), 50–100 µM BaCl_2_ and 1% PenStrep (SV30010; HyClone/Cytiva). In some experiments, recombinant GCAP1 was dissolved at 64.5 µM in 100 µL incubation solution/retina and remotely injected via thin tubing to minimize ambient perturbation, thereby achieving a final concentration of 3.2 µM at the photoreceptors; in these experiments, one retina from an animal acted as ‘treated’ while the other acted as ‘control’. Treated and control wells were alternated at every experiment. ERG records were analyzed with custom scripts. For each sequence, responses to the same flash strength were averaged and the following parameters extracted: Hill fit coefficient, saturating amplitude (S), half saturating flash strength (i_50_) and time-to-peak at i_50_ (TTP@i_50_). Parameter evolution during protein incubation was assessed after dual normalization: each retina relative to its own pre-treatment value and treated over control retina (*47*).

### Behavioral tests

All behavioral tests were performed at P270 in WT, E111V⁺/⁻ and E111V⁺/⁺ mice over a two-week period. Mice were habituated to the experimental room for at least 1 h before testing under dim light conditions; whiskers were trimmed the day before visual tests to minimize non-visual sensory inputs. All sessions were recorded with an infrared-sensitive camera and analyzed using ANY-maze v7.61 (Stoelting Co.).

Cognitive function (Novel Object Recognition, NOR), working memory (Y-maze spontaneous alternation), anxiety-like behavior (Elevated Plus Maze, EPM), and social interaction (Three-Chamber test) were assessed using standard protocols (*48, 49*). Briefly, in the NOR test, a discrimination index (DI = [novel − familiar] / total exploration time) was calculated as a measure of recognition memory 2 h after the training phase; animals with <20 s of total exploration time during either phase were excluded from the analysis. In the Y-maze, used to assess spatial working memory, spontaneous alternation (%) was calculated as the number of correct alternations divided by the total number of arm entries minus two, multiplied by 100. The OFT ( 45 × 45 cm arena, 10 min) was used to assess spontaneous exploratory locomotion and anxiety-like behavior by measuring total distance travelled, mean speed, and time spent in the center relative to the periphery of the arena. In the EPM, anxiety-like behavior was evaluated by quantifying the number of entries into and the time spent in the open arms. The three-chamber sociability test was used to assess sociability and social preference; following a 10-min habituation period, mice were allowed to freely explore the apparatus containing either an age- and sex-matched conspecific or a novel object.

Depth perception and visuospatial processing were evaluated with the Visual Cliff test and the Pole Descent Visual Cliff task (*50*). In the Visual Cliff test (25 × 55 cm transparent surface over a checkered floor, 8 min), time spent on each side and latency to first entry onto the shallow (“safe”) side were recorded. The Pole Descent Visual Cliff task (adapted from (*26, 50*)) consisted of 10 descending trials onto a 60 × 60 cm glass surface divided into one shallow quadrant and three cliff quadrants (30 cm depth); the percentage of correct (“safe”) landings was calculated. If, for three consecutive trials, a mouse did not exit the pole within three minutes, it was no further tested and excluded from the analysis.

### Transmission electron microscopy

The eyecups (enucleated as described for immunohistochemistry) were fixed in 2.5% glutaraldehyde and 2% paraformaldehyde solution for 2 h at 4°C, post-fixed in 1% OsO4 and 1.5% K4Fe(CN)6 for 2 h at 4°C, dehydrated in acetone and embedded in Epon 812 resin. Ultrathin (70 nm thick) sections were stained with Reynolds’ lead citrate for 1 min and observed with a Philips Morgagni transmission electron microscope operating at 80 kV and equipped with a Megaview III camera for digital image acquisition.

### Local field potential recordings and visual stimulation

Mice were initially anesthetized with 2% isoflurane in O_2_ (0.5 L/min), then placed into a stereotaxic apparatus. A small, custom-made plastic chamber was glued (Vetbond, St. Paul, MN, US) to the exposed skull. Animals were placed in a custom-made hammock, maintained under isoflurane anesthesia (1-2% in N_2_O/O_2_), and multiple single tungsten electrodes were inserted into a small craniotomy above the visual cortex. Once the electrodes were inserted, the chamber was filled with sterile agar and sealed with sterile bone wax. During recording sessions, animals were kept under isoflurane anesthesia (0.5 – 1% in O_2_). EEG and EKG were monitored throughout the experiments, and body temperature was maintained with a heating pad (in-house design).

Data were acquired using a 32-channel Scout recording system (Ripple, UT, USA). The local field potential (LFP) from multiple locations was bandpass-filtered from 0.1 Hz to 250 Hz and stored, along with spiking data, on a computer with at a 1 kHz sampling rate. The LFP signal was cut according to stimulus time stamps and averaged across trials for each recording location to calculate visually evoked potentials (VEP) (*51–54*). The spike signal was bandpass-filtered from 500 Hz to 7 kHz and stored on a computer hard drive at a 30 kHz sampling frequency. Spikes were sorted online in Trellis (Ripple, UT, USA) while performing visual stimulation. Visual stimuli were generated in MATLAB (Mathworks, USA) using Psychophysics Toolbox (*55*) and displayed on a gamma-corrected LCD monitor (35 inches, 100 Hz; 1920 x 1080 pixels; 52 cd/m^2^ mean luminance). Stimulus onset times were corrected for LCD monitor delay using a photodiode and microcontroller (in-house design) (*56*).

Vision was assessed using protocols published in our previous work (*52, 53, 56–59*). For recordings of visually evoked responses, cells were first tested with 100 repetitions of a 500 ms bright flash of light (300 cd/m^2^).

As the first step, evoked potentials across all layers were recorded, and the strongest response was used for comparisons between groups at the same cortical layer. The response amplitude of the VEP was calculated as the difference between the peak of the positive and negative components in the VEP wave. The maximum of the response was defined as the maximum of either the negative or positive peak.

### Statistics

Retinal morphology data were analyzed using GraphPad Prism v10.6.1 (GraphPad Software, San Diego, CA, USA). An ordinary one-way ANOVA with two-sided Dunnett’s multiple comparisons test was used to assess differences among groups; results are reported as mean ± SD. Variance homogeneity was evaluated using Bartlett’s test and Brown–Forsythe test. ERG data were analyzed using two-tailed Student’s independent samples t-test and are reported as mean ± SE; statistical comparisons were performed with KaleidaGraph v5.0.6 and JASP v0.95.4 (jasp-stats.org; RRID: SCR_015823). Behavioral data were analyzed by one-way ANOVA followed by two-sided Tukey’s multiple comparison test; results are reported as mean ± SE. Local field potential recordings were compared using two-tailed Mann–Whitney U-tests; offline data analysis and statistics were performed in MATLAB (MathWorks, Natick, MA, USA). For all the above analyses, exact *F* values, degrees of freedom, and *t* values are reported in the figure legends or supplementary tables. Differences were considered statistically significant at *P* < 0.05; significance levels are indicated as \**P* < 0.05, \*\**P* < 0.01, \*\*\**P* < 0.001.

For transcriptomic analyses, DEA was performed using the DESeq2 package v1.46.0 (*42*), which estimates variance-mean dependence in count data from high-throughput sequencing and assesses differential expression through a model based on the negative binomial distribution. Multiple testing correction was applied using the Benjamini–Hochberg false discovery rate (FDR) procedure throughout, including for DEA, GO enrichment, and GSEA. Genes and gene sets were considered statistically significant at FDR-adjusted *p* < 0.05; for differential expression, an additional effect-size threshold of |log₂FoldChange| > 1 was required. All transcriptomic analyses were performed in R v4.4.1.

## Data Availability

The RNA-seq raw data generated in this study have been deposited in the European Nucleotide Archive with project number PRJEB111775. The accession will be made publicly available upon publication. Source data underlying all graphs and figures (retinal morphology, ERG, behavioral, VEP, and immunofluorescence quantifications) are available from the corresponding authors upon reasonable request.

## Code Availability

All analyses used publicly available software (FastQC, Cutadapt, STAR, DESeq2, clusterProfiler, ggplot2; versions and parameters in Methods). The custom scripts implementing these described steps contain no novel algorithms and are available from the corresponding authors on request.

## ACKNOWLEDGEMENT

The authors thank Prof. Raffaella Mariotti for her support and training and Dr. Francesca Griggio for valuable assistance with preparation of RNA libraries. This work was supported by Retina Italia ODV (fellowship to A.A.), the Velux Foundation (Project No. 1410, to D.D.O. and L.C.), and by the Italian Ministry of University and Research under the National Recovery and Resilience Plan (NRRP) – Next Generation EU: the “Tuscany Health Ecosystem (THE)” project (CUP I53C22000780001, milestones 4.3.2, spoke 4, and 8.9.1, spoke 8) and the “A multiscale integrated approach to the study of the nervous system in health and disease (MNESYS)” project (CUP B33C22001060002, PE00000006, missione 4, componente 2, investimento 1.3). High-performance computing resources were provided through the “CONVECS” infrastructure, funded by the PR Veneto FESR 2021–2027 program (Priority 1, Specific Objective 1.1, Action 1.1.2). The Centro Piattaforme Tecnologiche of the University of Verona is acknowledged for access to spectroscopic, imaging, and genomic/transcriptomic platforms. Additional support was provided by the International Centre for Translational Eye Research (ICTER), carried out within the International Research Agendas program of the Foundation for Polish Science (FNP), co-financed by the European Union under the European Funds for Smart Economy 2021–2027 (FENG), and by TRIO-Vi CoE (No. 101136570). This work was also supported by the National Science Centre, Poland (grant no. 2022/47/B/NZ5/03023, to A.T.F.).

## AUTHOR CONTRIBUTION STATEMENT

A.A., G.D.C., A.B. and C.L. conducted morphology and immunohistochemistry/immunofluorescence experiments and analyses. A.A. conducted transcriptomic experiments. G.D.C, A.B. and V.M. produced the recombinant proteins. L.V. performed bioinformatic analyses under the supervision of G.M and with the contribution of V.M. B.C. conducted TEM experiments and analyzed data. G.T. and M.C. conducted behavioral experiments and analyzed data. K.S. performed VEP experiments and analyzed data under the supervision of A.F. S.A. and L.C designed and conducted ERG experiments. D.D.O. and L.C. conceived and designed the study. D.D.O. coordinated the research team. A.A. and D.D.O. wrote the manuscript, with contributions from all authors. All authors reviewed, edited, and approved the manuscript.

