## Supplementary Figures for "Retinal network dysfunction precedes structural degeneration in severe *GUCA1A* cone-rod dystrophy"

This file includes:

**Supplementary Figure S1.** Hematoxylin and eosin immunohistochemistry.

**Supplementary Figure S2**. Open field test.

**Supplementary Figure S3.** Behavioral analysis of cognition, sociability and anxiety-like behavior.

**Supplementary Figure S4.** Principal component analysis (PCA) of transcriptomic profiles from E111V$^{+/+}$ and WT mice at P30.

**Supplementary Figure S5.** Heatmap of the top differentially expressed genes between E111V$^{+/+}$ and WT mice at P30.

**Supplementary Figure S6.** Principal component analysis (PCA) of transcriptomic profiles from E111V$^{+/+}$ and WT mice at P270.

**Supplementary Figure S7.** Heatmap of the top differentially expressed genes between E111V$^{+/+}$ and WT mice at P270.

**Supplementary Figure S8.** Dot plot summarizing Gene Ontology (GO) enrichment analysis across the four transcriptomic comparisons.

**Supplementary Figure S9.** Dot plot of Gene Set Enrichment Analysis (GSEA) results for the comparison E111V$^{+/+}$ mice at P270 *versus* P30.

**Supplementary Figure S10.** Dot plot of Gene Set Enrichment Analysis (GSEA) results for the comparison E111V$^{+/+}$ *versus* WT mice at P30.

**Supplementary Figure S11.** Dot plot of Gene Set Enrichment Analysis (GSEA) results for the comparison E111V$^{+/+}$ *versus* WT mice at P270.

**Supplementary Movie 1.** Pole Descent Visual Cliff Task in WT mice. Representative movie of a pole visual cliff experiment with a WT mouse descending a vertical pole onto a 60 × 60 cm glass platform (one shallow quadrant, three 30 cm-drop cliff quadrants), used to assess depth perception and visuospatial processing. The movie spans a fraction of the 330s of real time measurement at 1× playback speed.

**Supplementary Movie 2.** Pole Descent Visual Cliff Task in E111V$^{+/+}$ mice. Representative movie of a pole visual cliff experiment with an E111V$^{+/+}$ mouse descending a vertical pole onto a 60 × 60 cm glass platform (one shallow quadrant, three 30 cm-drop cliff quadrants), used to assess depth perception and visuospatial processing. The movie spans a fraction of the 330s of real time measurement at 1× playback speed.

**Supplementary Data 1**. Definition of targeted biological pathways analyzed in this study. For each pathway, the table reports the associated curated functional categories used for targeted analyses. Biological pathways were color-coded to facilitate visual comparison of the enrichment analysis presented in Supplementary Data 7 and Supplementary Data 8 as follows: Retina function and development (blue), Synapse (orange), Death (gray), Inflammation (yellow), Metabolism (green), Cytoskeleton (pink). The targeted analyses were intended as hypothesis-guided exploratory analyses complementary to the unbiased GO and GSEA approaches.

**Supplementary Data 2.** Summary of sequencing data quality metrics before and after trimming all samples. For each analysis ID, genotype (GT; MUT=E111V$^{+/+}$ WT=wild type), and time point, the table reports the total number of raw sequences, read length, GC content prior to trimming, as well as the number and percentage of reads retained after trimming.

**Supplementary Data 3.** Differential gene expression analysis between E111V$^{+/+}$ and WT samples at timepoint P30 based on RNA-seq data. For each gene, the table reports the mean normalized expression level (baseMean), log2 fold change between E111V$^{+/+}$ and WT, standard error of the log2 fold change (lfcSE), test statistic, raw p-value, and adjusted p-value (padj).

**Supplementary Data 4.** Differential gene expression analysis between E111V$^{+/+}$ and WT samples at timepoint P270 based on RNA-seq data. For each gene, the table reports the mean normalized expression level (baseMean), log2 fold change between E111V$^{+/+}$ and WT, standard error of the log2 fold change (lfcSE), test statistic, raw p-value, and adjusted p-value (padj).

**Supplementary Data 5.** Differential gene expression analysis between P270 vs P30 in E111V$^{+/+}$ samples based on RNA-seq data. For each gene, the table reports the mean normalized expression level (baseMean), log2 fold change between P270 and P30, standard error of the log2 fold change (lfcSE), test statistic, raw p-value, and adjusted p-value (padj).

**Supplementary Data 6.** Differential gene expression analysis between P270 vs P30 in WT samples based on RNA-seq data. For each gene, the table reports the mean normalized expression level (baseMean), log2 fold change between P270 and P30, standard error of the log2 fold change (lfcSE), test statistic, raw p-value, and adjusted p-value (padj).

**Supplementary Data 7.** Gene Ontology (GO) enrichment analysis of differentially expressed genes. For each comparison and GO term, the table reports the GO identifier, term description, gene ratio, background ratio, rich factor, fold enrichment, z-score, p-value, adjusted p-value, q-value, the list of associated genes, and the number of genes contributing to the enrichment (Count). GO terms were color-coded according to the biological pathway they belong to, as defined in Supplementary Data 1: Retina function and development (blue), Synapse (orange), Death (gray), Inflammation (yellow), Metabolism (green), Cytoskeleton (pink).

**Supplementary Data 8.** Gene Set Enrichment Analysis (GSEA) results. For each comparison and gene set, the table reports the gene set identifier and description, set size, enrichment score (ES), normalized enrichment score (NES), p-value, adjusted p-value, q-value, rank at maximum enrichment, leading-edge subset, and the core enriched genes driving the enrichment. GO terms were color-coded according to the biological pathway they belong to, as defined in Supplementary Data 1: Retina function and development (blue), Synapse (orange), Death (gray), Inflammation (yellow), Metabolism (green), Cytoskeleton (pink).

### *Supplementary Figures*

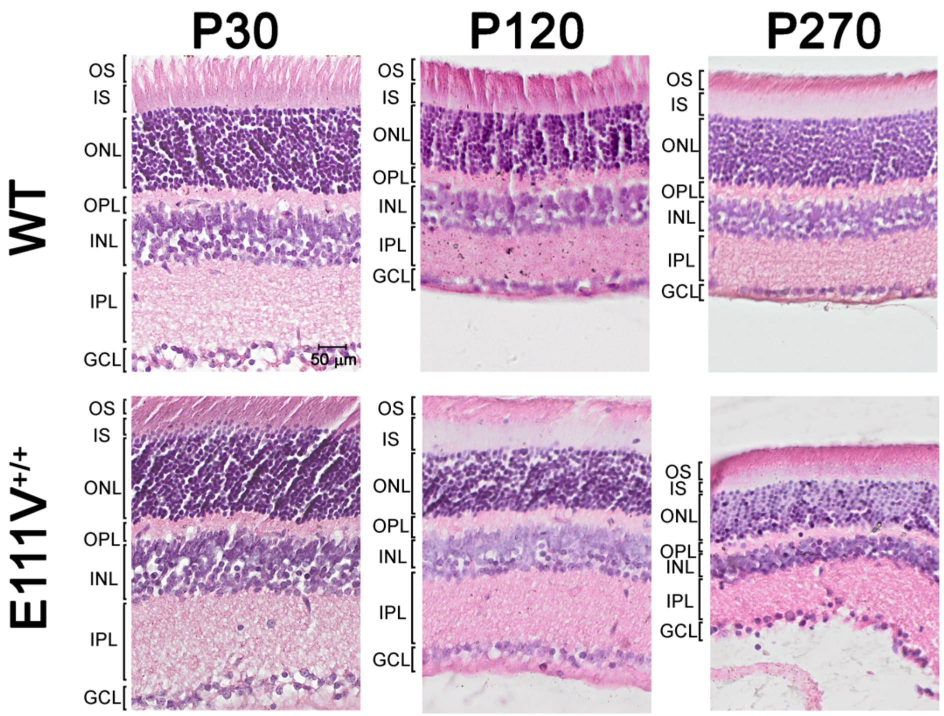


**Supplementary Figure S1. Hematoxylin and eosin immunohistochemistry.** Representative images from six animals (n=6) per genotype wild-type (WT) and E111V$^{+/+}$ at various time points from P30 to P270.

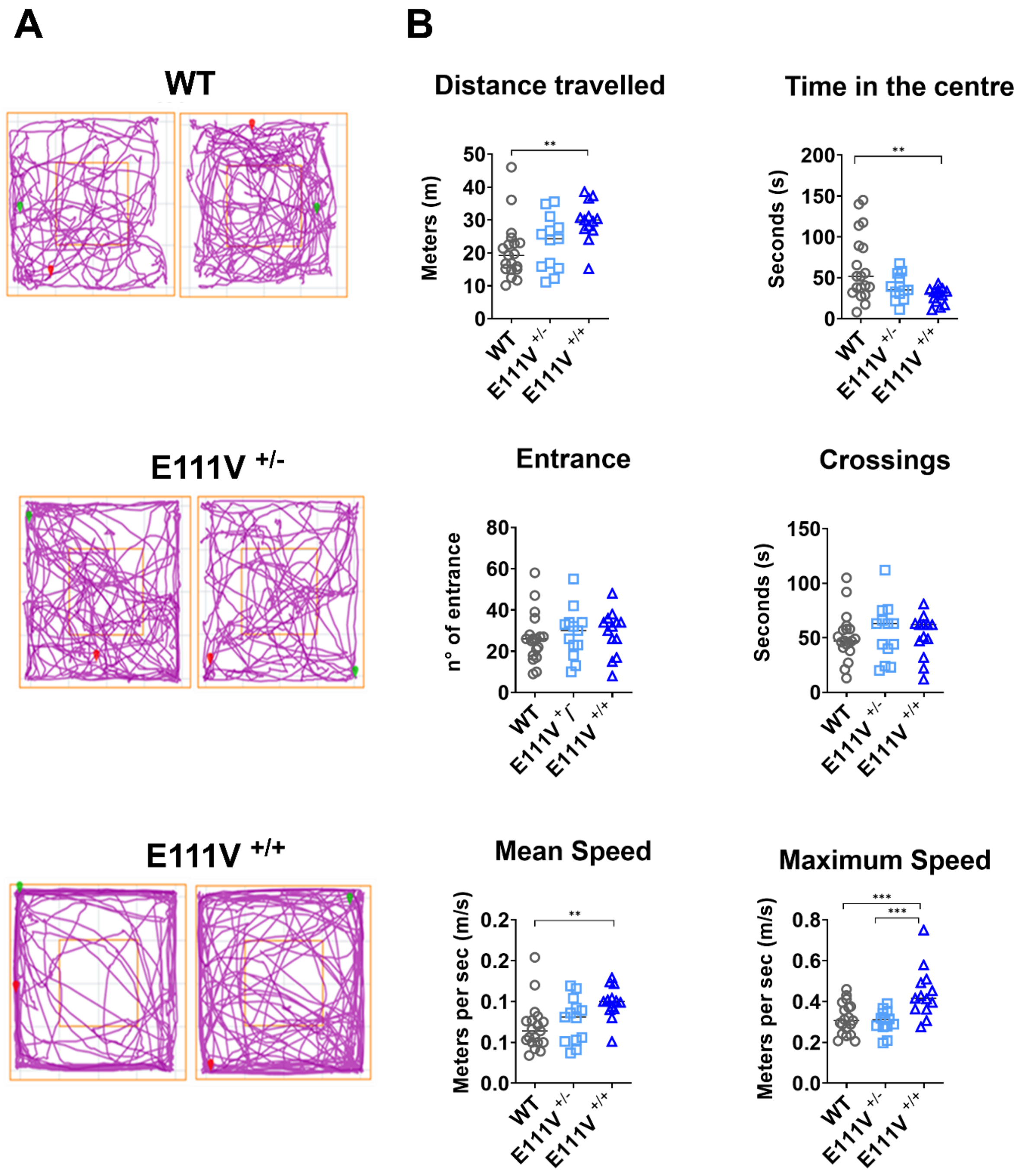


**Supplementary Figure S2. Open field test.** A) Two representative graphs per group showing the activity of the mice in the open field test. B) Total distance travelled in the arena, time spent in the central zone, number of entries into the arena, time spent crossing from the outer to inner of arena, mean speed and maximum speed. WT (n=19), E111V$^{+/-}$ (n=13) and E111V$^{+/+}$ (n=13) at P270.

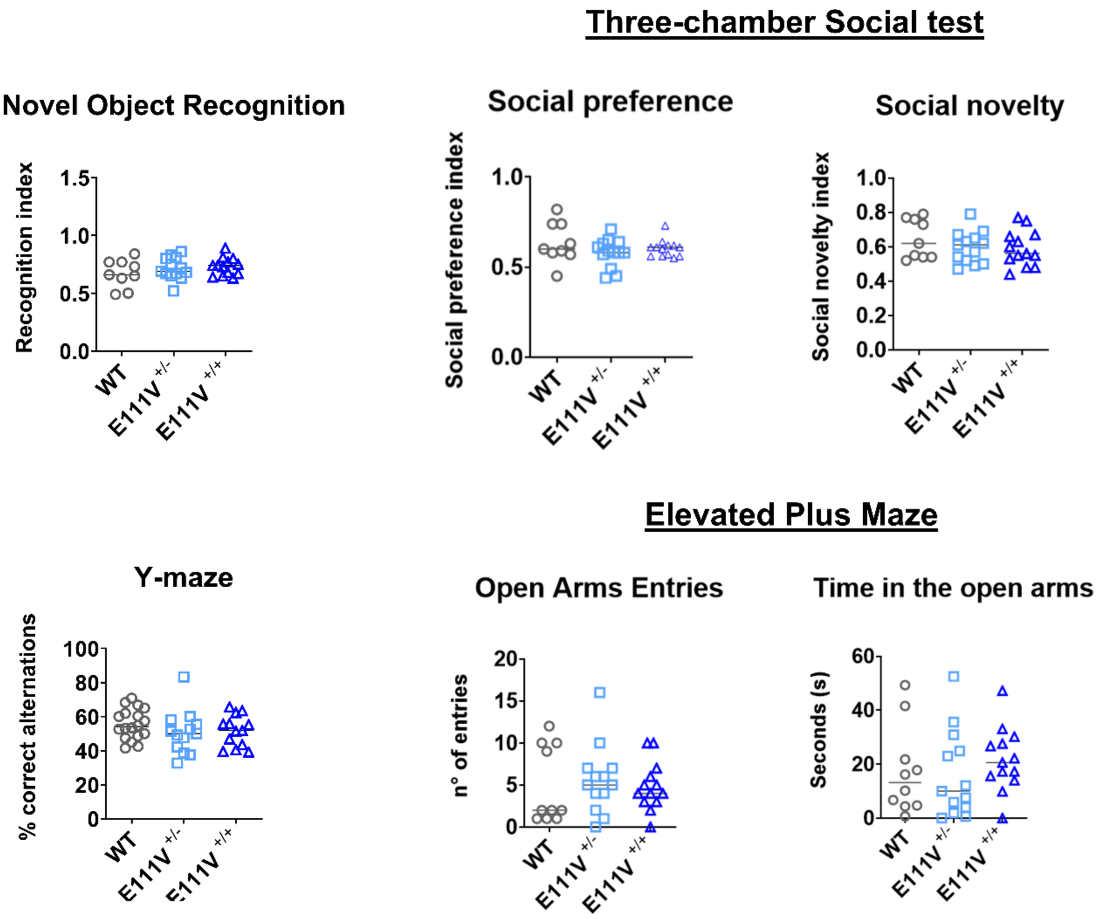


**Supplementary Figure S3. Behavioral analysis of cognition, sociability and anxiety-like behavior.** Novel Object Recognition (NOR) recognition index, Three-Chamber Social Test social preference and social novelty indices, Y-maze spontaneous alternation performance, and Elevated Plus Maze (EPM) open-arm entries and time spent in the open arms in WT (n=9), E111V$^{+/-}$ (n=13) and E111V$^{+/+}$ (n=13) mice at P270.

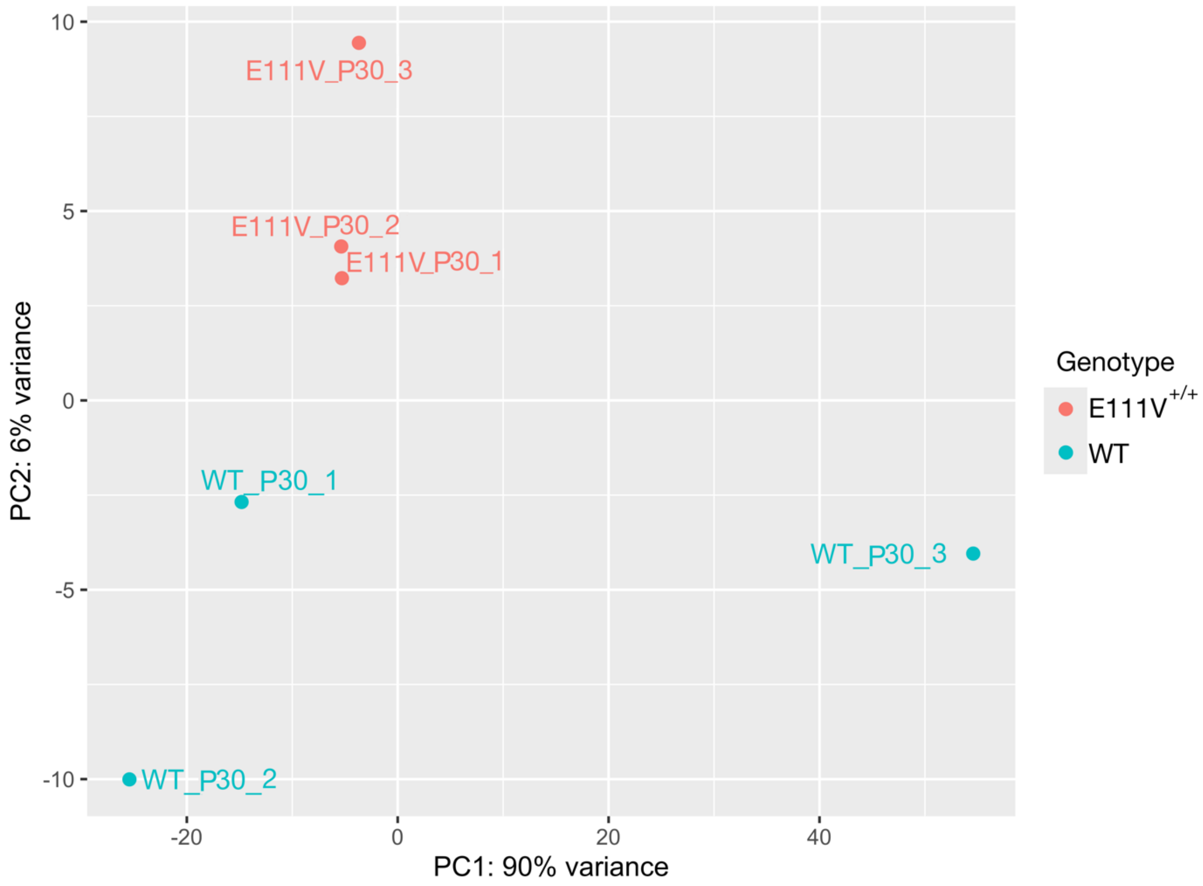


**Supplementary Figure S4. Principal component analysis (PCA) of transcriptomic profiles from E111V$^{+/+}$ and WT mice at P30.** The PCA was performed on normalized gene expression data, illustrating the global transcriptional differences between genotypes at this time point.

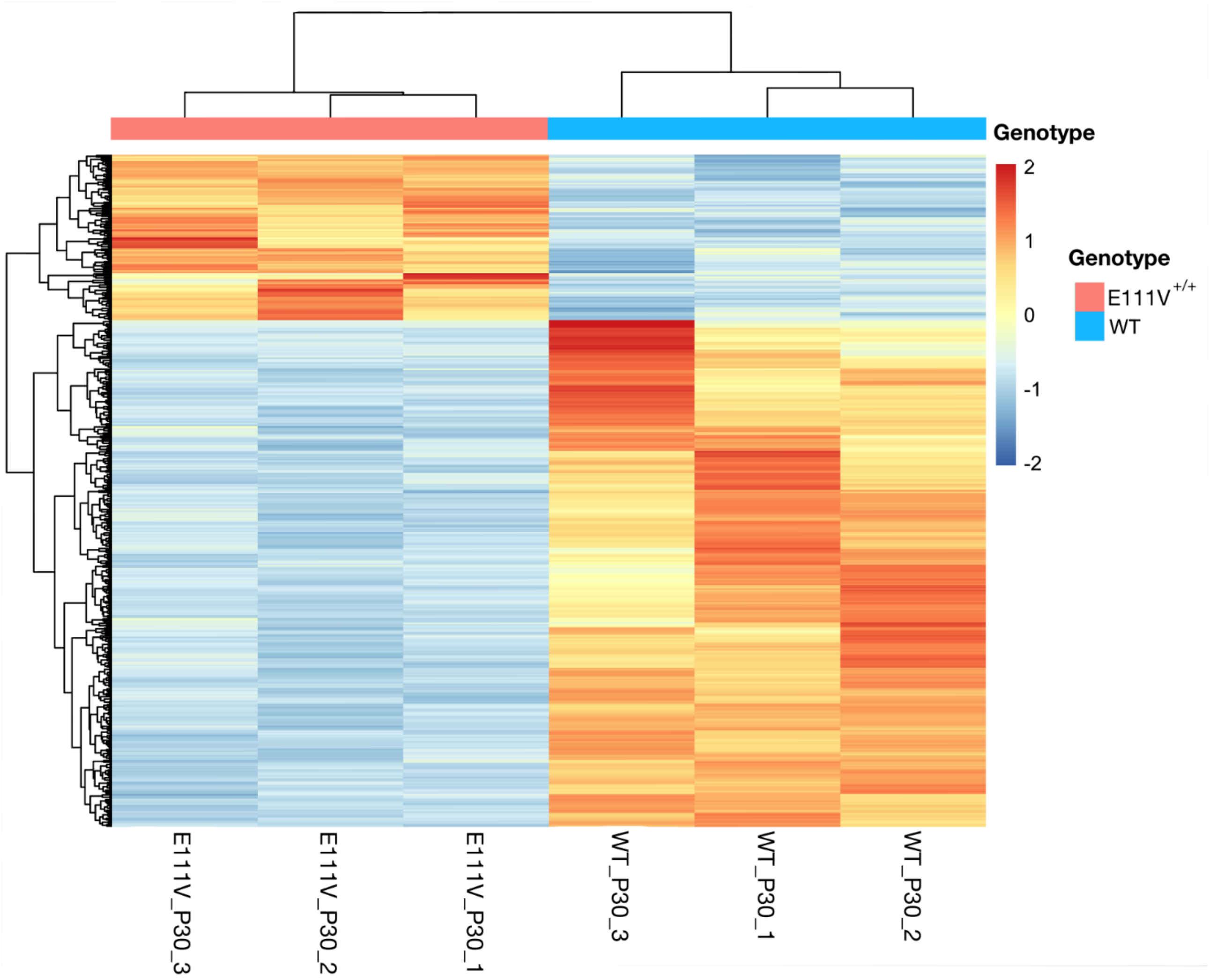


**Supplementary Figure S5. Heatmap of the top differentially expressed genes between E111V$^{+/+}$ and WT mice at P30.** Rows represent genes and columns represent individual samples. Gene expression values were normalized and scaled by gene to highlight relative expression differences across samples.

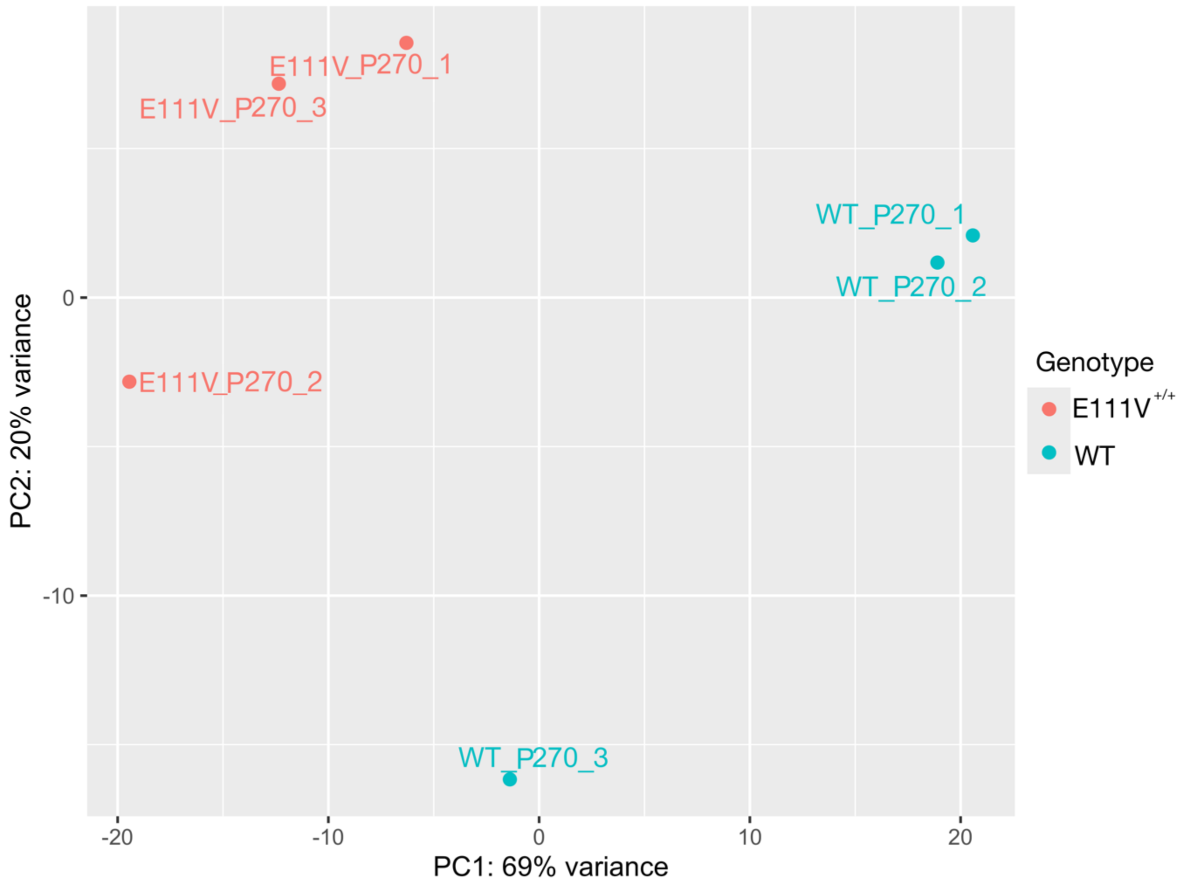


**Supplementary Figure S6. Principal component analysis (PCA) of transcriptomic profiles from E111V$^{+/+}$ and WT mice at P270.** The PCA was performed on normalized gene expression data, illustrating the global transcriptional differences between genotypes at this time point.

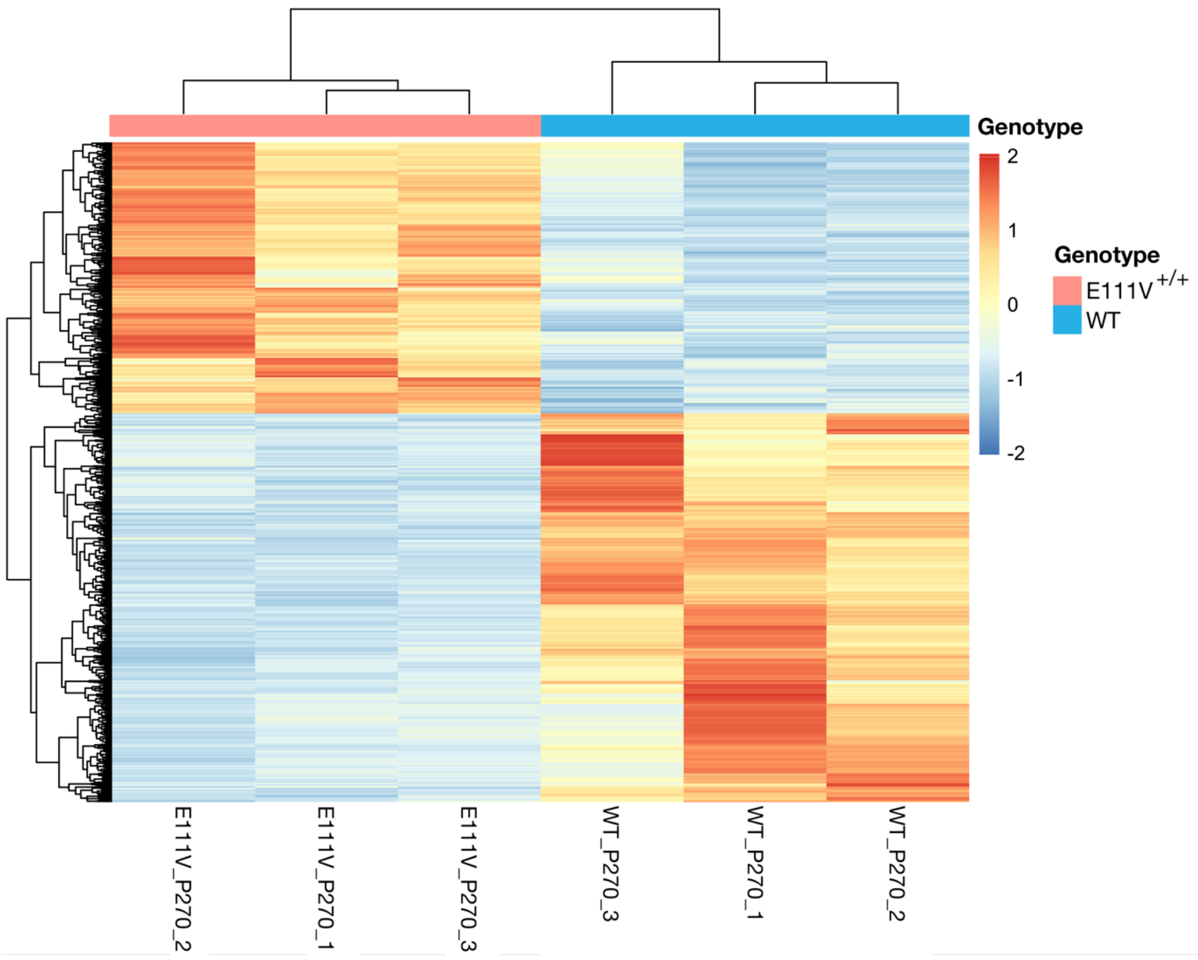


**Supplementary Figure S7. Heatmap of the top differentially expressed genes between E111V$^{+/+}$ and WT mice at P270.** Rows represent genes and columns represent individual samples. Gene expression values were normalized and scaled by gene to highlight relative expression differences across samples.

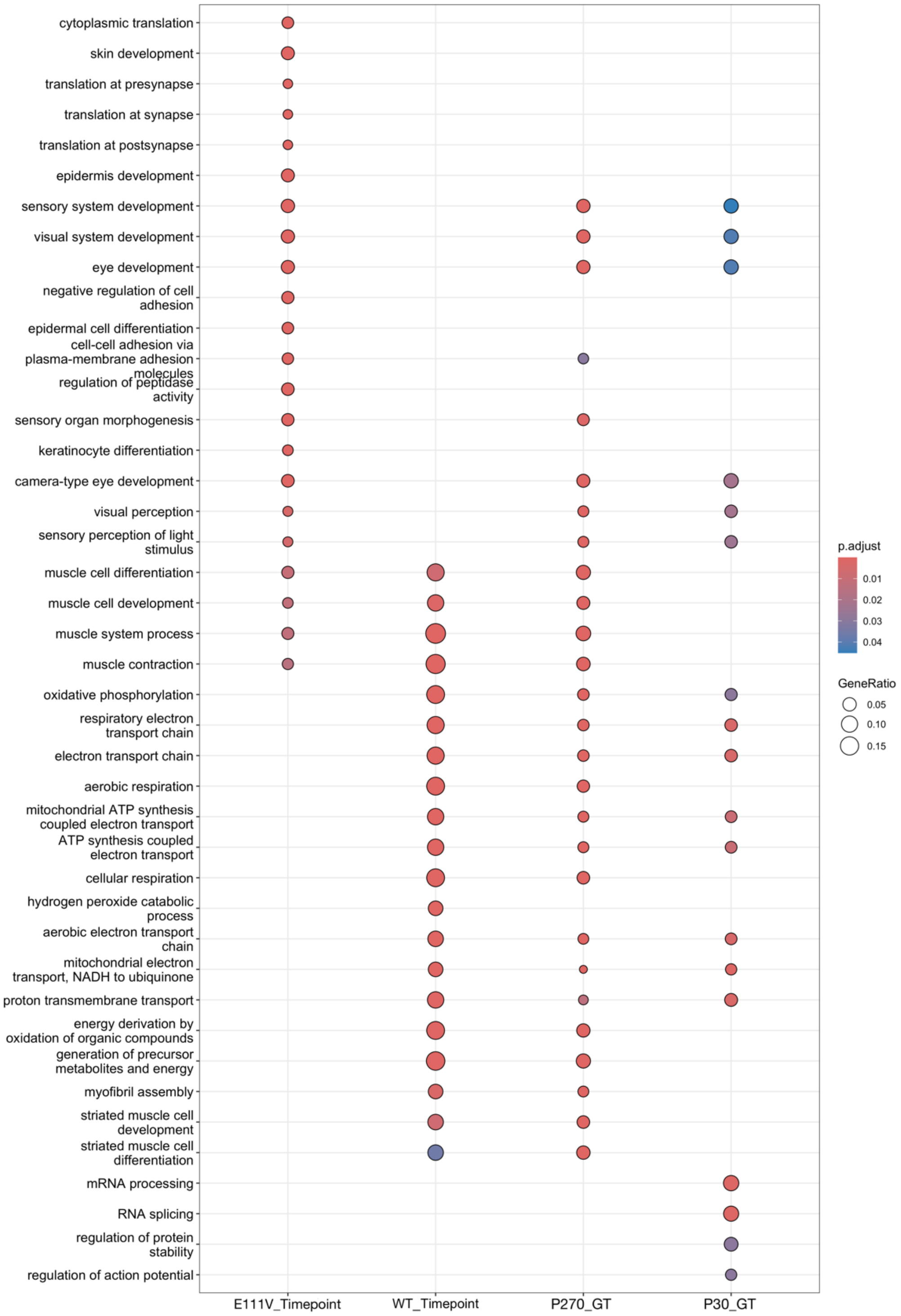


**Supplementary Figure S8. Dot plot summarizing Gene Ontology (GO) enrichment analysis across the four transcriptomic comparisons.** Each dot represents an enriched GO term, with color indicating the adjusted p-value (blue to red) and dot size representing the GeneRatio. The four contrasts are defined as follows: E111V_Timepoint = E111V$^{+/+}$ mice at P270 versus P30; WT_Timepoint = WT mice at P270 versus P30; P270_GT = E111V$^{+/+}$ versus WT mice at P270; and P30_GT = E111V$^{+/+}$ versus WT mice at P30.

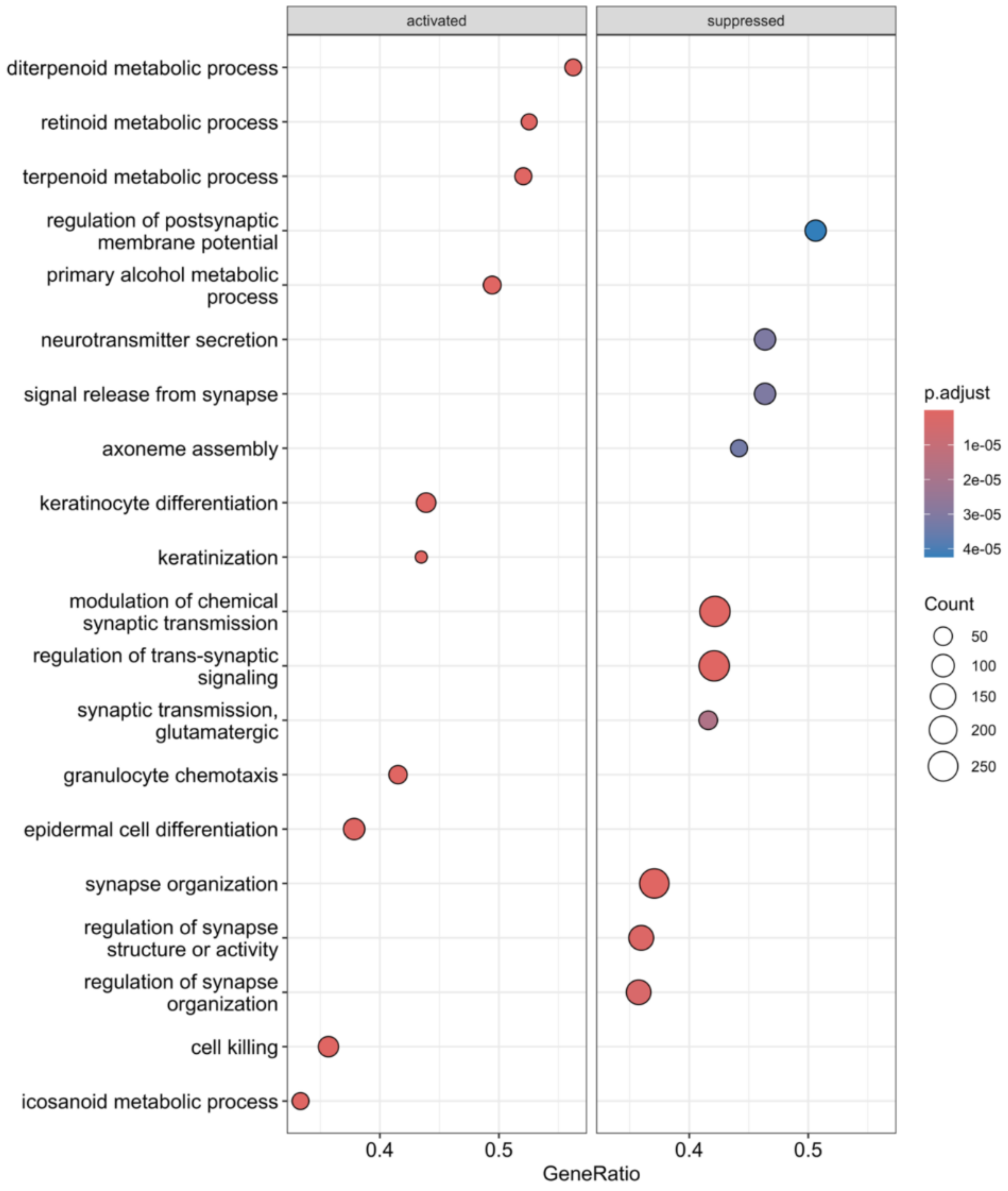


**Supplementary Figure S9. Dot plot of Gene Set Enrichment Analysis (GSEA) results for the comparison E111V$^{+/+}$ mice at P270 versus P30.** Activated and suppressed gene sets are shown, with dot color indicating the adjusted p-value (blue to red) and dot size representing the GeneRatio. Gene sets with positive and negative enrichment correspond to pathways activated and suppressed at P270 relative to P30, respectively.

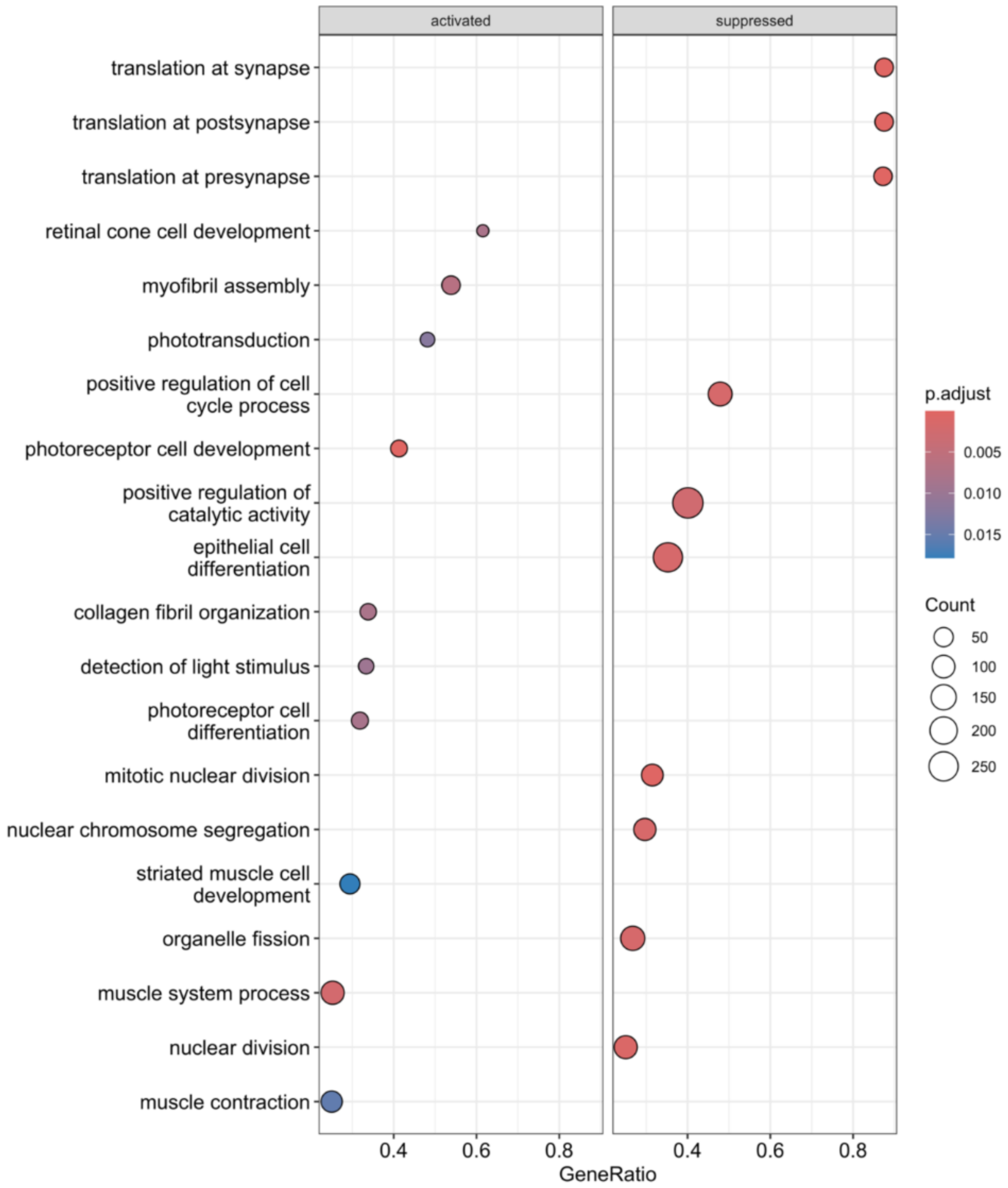


**Supplementary Figure S10. Dot plot of Gene Set Enrichment Analysis (GSEA) results for the comparison E111V$^{+/+}$ vs WT mice at P30.** Activated and suppressed gene sets are shown, with dot color indicating the adjusted p-value (blue to red) and dot size representing the GeneRatio. Gene sets with positive and negative enrichment correspond to pathways activated and suppressed in E111V$^{+/+}$ relative to WT, respectively.

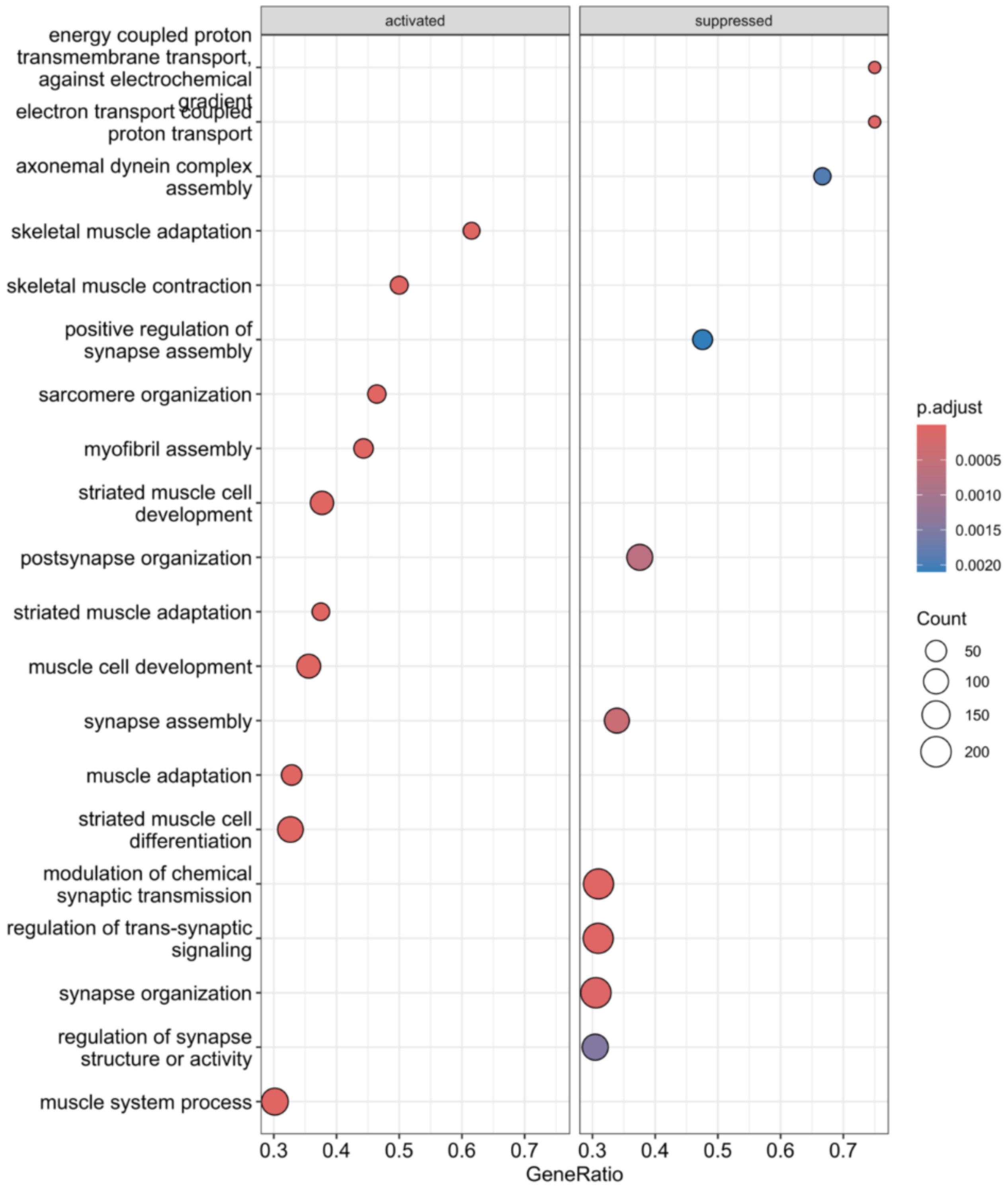


**Supplementary Figure S11. Dot plot of Gene Set Enrichment Analysis (GSEA) results for the comparison E111V$^{+/+}$ vs WT mice at P270.** Activated and suppressed gene sets are shown, with dot color indicating the adjusted p-value (blue to red) and dot size representing the GeneRatio. Gene sets with positive and negative enrichment correspond to pathways activated and suppressed in E111V$^{+/+}$ relative to WT, respectively.
